# A cell circuit approach to dissect fibroblast-macrophage interactions in the tumor microenvironment

**DOI:** 10.1101/2022.11.17.516850

**Authors:** Shimrit Mayer, Tomer Milo, Achinoam Isaacson, Coral Halperin, Shoval Miyara, Yaniv Stein, Meirav Pevsner-Fischer, Eldad Tzahor, Uri Alon, Ruth Scherz-Shouval

## Abstract

The tumor microenvironment (TME) is composed of various nonmalignant cell types that interact with each other and with cancer cells, impacting all aspects of cancer biology. The TME is complex and heterogeneous, and thus simplifying systems and concepts are needed. Here we provide a tractable experimental system and powerful mathematical circuit concepts to identify the main molecular interactions that govern the composition of the TME. We focus on two major components of the TME - cancer associated fibroblasts (CAFs) and tumor associated macrophages (TAMs), define their interactions and verify our predictions in mouse and human breast cancer. We measure the population dynamics starting from many initial conditions of co-cultures of macrophages and organ-derived fibroblasts from mammary, lung, and fat, and explore the effects of cancer-conditioned medium on the circuits. We define the circuits and their inferred parameters from the data using a mathematical approach, and quantitatively compare the cell circuits in each condition. We find that while the homeostatic steady-states are similar between the organs, the cancer-conditioned medium profoundly changes the circuit. Fibroblasts in all contexts depend on autocrine secretion of growth factors whereas macrophages are more dependent on external cues, including paracrine growth factors secreted from fibroblasts and cancer cells. Transcriptional profiling reveals the molecular underpinnings of the cell circuit interactions and the primacy of the fibroblast autocrine loop. The same fibroblast growth factors are shared by the co-cultures and mouse and human breast cancer. The cell circuit approach thus provides a quantitative account of cell interactions in the cancer microenvironment.

## Introduction

Tumors are complex ecosystems in which cancer cells interact with diverse non-malignant cells of the tumor microenvironment (TME). It is through these interactions that tumors progress and metastasize, and these interactions impact all aspects of cancer biology ^1–3^. Major components in the tumor microenvironments (TME) of most carcinomas are fibroblasts and macrophages, known as cancer associated fibroblasts (CAFs) and tumor associated macrophages (TAMs) ^4,5^. Defining the inherent principles and molecular signals of CAF and TAM interactions is critical for our understanding of the TME and for finding ways to improve cancer therapy.

Fibroblasts and macrophages are basic tissue components of every organ in the body, and key regulators of organ homeostasis ^6,7^. In healthy tissues, fibroblasts produce the extracellular matrix (ECM) that gives structure to the organ and limits the proliferation and differentiation of epithelial cells ^8,9^. Macrophages serve as sentinels that detect stress signals and phagocytose invading pathogens, apoptotic cells and ECM degradation products ^6^. Upon injury or inflammation, circulating monocytes infiltrate the organ and differentiate into macrophages. These bone-marrow-derived macrophages (BMDMs) ^10^ work together with the resident macrophages and fibroblasts. The resident fibroblasts transition into *myofibroblasts* that produce copious ECM, which is further remodeled by the BMDMs in a reciprocal process of wound healing. Both in normal homeostasis and following injury, fibroblasts have additional roles beyond ECM production, influencing epithelial stem cell behavior, promoting angiogenesis and coordinating immune function through production of chemokines and cytokines ^11,12^.

In severely injured or chronically inflamed tissues, fibroblasts produce excessive ECM, and macrophages remodel the ECM through production of ECM-modifying enzymes such as metalloproteases ^8,13^. The excessive production and remodeling of ECM may result in fibrosis, in which tissue is replaced with scar. Eventually, fibrosis can lead to cancer ^14,15^. Fibroblasts in cancer are rewired to become protumorigenic CAFs that support tumor progression, invasion, and metastasis by secreting cytokines, chemokines, extracellular matrix components and growth factors ^11,12^. CAFs also promote the recruitment of monocytes from the bone-marrow and their differentiation into TAMs, which then stimulate angiogenesis, enhance tumor cell migration and invasion, and suppress antitumor immunity ^14^.

Fibroblasts and macrophages are transcriptionally and phenotypically heterogeneous. One facet of this heterogeneity is that different organs have different characteristic populations of resident fibroblasts. Advances in single-cell RNA sequencing and imaging technologies defined multiple subpopulations of fibroblasts in healthy, and even more so in diseased tissues ^16^. These subpopulations have distinct functions related to ECM production, adhesion and immune regulation ^17–19^. In many cases these subpopulations represent dynamic cell states induced by changes in microenvironmental conditions ^18^. Infiltrating macrophages come from a shared monocyte origin, and can switch between pro-inflammatory and anti-inflammatory states, contributing to phenotypic plasticity of diseased tissues ^20,21^.

Heterogeneous populations of fibroblasts and macrophages engage in complex cell-cell interactions. To understand these interactions, simplifying concepts are essential. One such concept is the cell circuit ^22^. Cell circuits describe the dynamics of cell numbers for several cell types according to the growth factors they exchange and constraints imposed by their environment. Recently, a prototype circuit model for fibroblast-macrophage interactions in tissue homeostasis was defined using *ex-vivo* co-cultures of mouse embryonic fibroblasts (MEFs) with BMDMs ^22,23^. Fibroblasts, however, are heterogeneous and evolve to perform organ-specific tasks ^11,16^. In tumors, fibroblasts evolve and diverge into distinct subpopulations with distinct functions such as immune-modulation, ECM production and antigen-presentation ^12,18,24,25^. It remains unknown what the cell circuits are in the cancer microenvironment, and how they differ from the normal cell circuits in different organs.

Here we address these questions by developing a cell circuit approach that combines experimental co-culture and mathematical modeling to infer circuit parameters, such as the autocrine and paracrine interactions between multiple cell types, and compare between different cell circuits. We apply this approach to co-cultures of BMDMs with fibroblasts from three different organs - mammary, lung, and mesometrial fat - and explore the effects of cancer-conditioned medium on the circuits. We find that the homeostatic steady states are similar between the organs, but changes in growth conditions from normal to cancer-conditioned medium profoundly alter the circuit interactions. In all contexts, fibroblasts support their own growth by an autocrine loop of growth factors, whereas macrophages are more dependent on external growth factors secreted by fibroblasts and cancer cells. RNA sequencing of the co-cultures supports the inferred circuit interactions and their relative strengths, and highlights potential growth factors driving these interactions. Comparative transcriptomic analysis of mouse and human breast cancer reveals that the fibroblast autocrine loop is the strongest interaction in all circuits, and that the growth factors that comprise it are shared by the circuits found in the *in-vitro-*simulated cancer microenvironment and the microenvironment of mouse and human breast cancer. Together, our findings establish principles of cell circuit design, and provide a quantitative approach to model cell interactions in an organ- and disease-specific manner.

## Results

### Experimental phase portraits show multiple steady-states for macrophage-fibroblast circuits which are similar across organs, but perturbed in cancer-conditioned medium

To define cell circuits in different organs and cancer contexts, we established a co-culture assay for cell population dynamics. As a baseline, we used fibroblasts from the mammary fat pad and later we compared them with fibroblasts from fat and lung to explore organ context. We then compared growth in control medium to breast cancer-conditioned medium (CM), to explore tumor context.

We co-cultured BMDMs with tissue-resident fibroblasts from the mammary fat pad of syngeneic BALB/c mice using a previously described approach for mouse embryonic fibroblasts ^22^. Cell growth from different initial concentrations of fibroblasts and BMDMs was tracked by flow cytometry after 3 or 7 days of co-culture (Figure S1A), and plotted in a *phase portrait*. The phase portrait has two axes: the fibroblast (X-axis) and macrophage (Y-axis) cell counts. Arrows (vectors) on the phase portrait indicate how cell counts change from day 3 to day 7 (Figure 1A).

**Figure 1:**
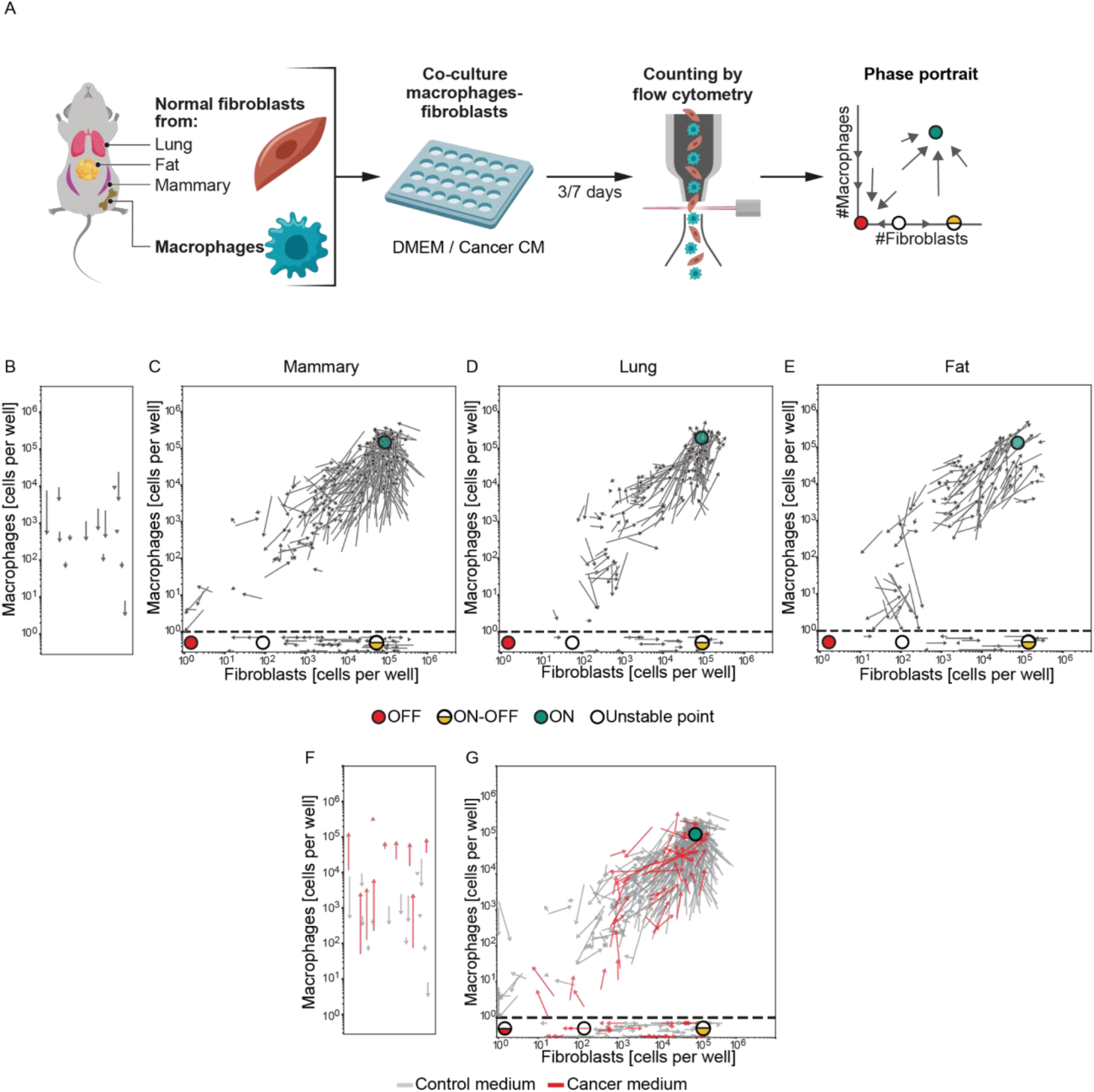
Phase portraits of fibroblasts and macrophages are similar in different organ contexts, but perturbed by cancer-conditioned medium. **A**. Illustration of the experimental procedure. Bone-marrow derived macrophages and fibroblasts from the indicated organs were isolated from mice, co-cultured at different ratios for 3 or 7 days in control or cancer-conditioned medium (CM), and then counted by flow cytometry. **B-G**. Experimental phase portraits of macrophage-fibroblast dynamics *in-vitro*. Arrow tails represent cell counts at day 3 of co-culture, and arrowheads represent cell counts at day 7 (starting from the same initial cell concentration). **B**. Mono-cultured macrophages are presented. **C-E**. Mammary, lung, and fat fibroblasts co-cultured with macrophages are presented above the horizontal dashed lines. Mono-cultured fibroblasts from these organs are presented below the dashed lines of these plots. Fixed points are denoted as follows: The ‘‘ON’’ state: green dot; ‘‘OFF’’ state: red dot; unstable point: white dot; ‘‘ON-OFF’’ state: half-yellow dot. Data are combined from at least three independent experiments, with the following total number of biological replicates: macrophages only: n=5; mammary: n=24; lung: n=16; fat: n=10. **F**. Experimental phase portrait of macrophages grown in mono-culture in the presence of conditioned media (CM) derived from 4T1 breast cancer cells (red arrows), overlayed on the control phase portrait presented in Figure 1B (gray arrows). Data are combined from 3 independent experiments (performed in parallel to control media cultures); n=10 biological replicates. **G**. Experimental phase portrait of macrophage-mammary fibroblast dynamics following *in-vitro* co-culture with 4T1 CM (red arrows), overlayed on the control mammary phase portrait presented in Figure 1C (gray arrows). Mammary fibroblasts co-cultured with macrophages are presented above the horizontal dashed line; mono-cultured fibroblasts are presented below the dashed lines of the plot. Data is combined from 5 independent experiments (performed in parallel to control media co-cultures); n=12 biological replicates for the cancer CM.

Macrophages in mono-culture could not promote their own growth under these experimental conditions (Figure 1B), but co-culture with fibroblasts supported macrophage growth, and revealed dynamic interactions (Figure 1C).

The phase portrait shows several points of interest that characterize the dynamic system, called fixed points, in which cell counts remain approximately constant ^22,26^. One fixed point is the ON state (Figure 1C, green dot) with high numbers of fibroblasts and macrophages. In this state, macrophages and fibroblasts continually turn over as indicated by EdU incorporation measurements (Figure S1B-C; see Methods), but maintain their numbers in a dynamic steady state. *In vivo*, this state may occur following injury, as fibroblasts and macrophages are recruited to repair wounded tissue. If the injury is not resolved, fibroblasts and macrophages maintain high numbers in a state of chronic inflammation and fibrosis ^10,23^.

When fibroblasts grow alone their growth dynamics depend on their initial numbers. Below a critical threshold, which is an unstable fixed point (Figure 1C, white dot), fibroblast numbers decline to zero. This represents the healthy state of a tissue, and it is the expected outcome of successful resolution of acute injury or acute inflammation. Above this threshold, fibroblasts are able to maintain themselves, and their numbers rise to a fixed point called the ON-OFF state (fibroblasts are ON, macrophages are OFF, Figure 1C, half-yellow dot). Fibroblasts at this fixed point continually turn over in a dynamic steady-state, as indicated by EdU incorporation (Figure S1B). In physiological terms, a state in which fibroblasts maintain high numbers in the absence of macrophages is referred to as ‘cold fibrosis’ ^23^, and it is distinct from the ON state, which has macrophages instead.

When macrophages are added to the ON-OFF state, the macrophage number increases and converges to the ON state (Figure 1C, green dot). Thus the ON-OFF fixed point is semi-stable, and the cold fibrosis state leads to ‘hot’ fibrosis.

An additional fixed point is at zero cells, denoted as the OFF state (Figure 1C, red dot). The phase portrait indicates that this state is reached when initial fibroblast and macrophage concentrations are low. Physiologically, as mentioned above, this can be viewed as a healing state, in which myofibroblasts and inflammatory macrophages are no longer present.

We tested the robustness of the phase portrait assay in several ways (Figure S1D-G). Biological replicates of the experiment gave rise to similar phase portraits (Figure S1D). Phase portraits derived from cell counts at days 7, 14, and 21 showed a qualitatively similar convergence towards the “ON” and “ON-OFF” states found for cell counts derived from days 3 and 7 (Figure S1E), suggesting approximate temporal invariance. We repeated the analysis using a different mouse strain, C57BL/6, and obtained a similar phase portrait (Figure S1F). To accurately measure growth dynamics at very low initial cell concentrations (several cells per well) we scaled the experiment up from 96-well plates to 6-well plates, which have a 30-fold larger area. We accordingly multiplied cell numbers measured in 96-wells by a factor of 30. We tested the validity of this approach by measuring overlapping regions of similar effective concentrations in 96 and 6 wells, and found qualitative agreement (Figure S1G, red vs. gray arrows).

We further asked whether addition of macrophages to an on-going culture of fibroblasts - simulating the infiltration of BMDMs into a tissue populated by resident fibroblasts - would yield similar interaction dynamics compared to those observed by simultaneous plating of both cell types. We observed similar convergence towards the ON state when macrophages were either added to the cultures 3 days after the initial plating of fibroblasts, or simultaneously plated with fibroblasts, suggesting that the interaction dynamics are independent of this variable (Figure S1G, pink arrows). This finding further supports the conclusion that the ON-OFF state is semi-stable.

Next, we used this assay to test the effect of organ context on the phase portrait. We sought to understand whether resident fibroblasts from different organs have similar or different interaction circuits with infiltrating macrophages. For this purpose, we isolated fibroblasts from two additional organs, lung and mesometrial fat. We determined the phase portrait for these fibroblasts grown with BMDMs (Figure 1D-E), and compared them to the phase portrait from mammary fibroblasts.

The phase portraits from all three organs showed similar fixed point structures and robustness criteria (Figure 1C-E, Figure S1D-G, and Figure S2A-F). The three portraits had an OFF state (red dots), an unstable fixed point located on the x axis (white dots), an ON-OFF state (half-yellow dots) in which fibroblasts maintain their own numbers, and an ON state (green dots) in which fibroblasts and macrophages support each other in a dynamic steady state.

The position of the fixed points was similar in all three organ contexts. The ON state was composed of ~10^5^ fibroblasts and ~10^5^ macrophages (Figure 1C-E), and the ON-OFF state had ~10^5^ fibroblasts in all three organs. Additionally, the fixed points showed similar cell sizes measured by microscopy (Figure S2G-H). These findings suggest that the homeostatic steady-state concentrations of fibroblasts and macrophages are similar across organ contexts.

We next asked how the circuits might be affected by pathological conditions. In particular, cancer cells rewire their microenvironment, with effects on fibroblasts and macrophages ^11,12,27^. To test this, we used cancer conditioned medium (CM) in the co-culture system and measured changes in the phase portraits of fibroblasts and macrophages, using the mammary fat pad as a baseline system.

We grew 4T1 triple-negative breast cancer cells syngeneic to the fibroblasts and BMDMs, and collected their CM at 48 hours. We added the cancer CM to co-cultures of mammary fibroblasts and BMDMs, and obtained the experimental phase portrait (Figure 1F-G; red arrows).

In the presence of cancer CM (Figure 1F-G) the phase portrait was very different from control medium (Figure 1C). Macrophages in cancer CM were able to grow in the absence of fibroblasts (Figure 1F), in contrast to control media, in which their growth was found dependent on fibroblasts (Figure 1B). This may relate to the composition of 4T1-conditioned medium which contains factors that regulate macrophage proliferation ^28^. As a result, the OFF state, which was stable in the control medium, becomes semi-stable in the presence of cancer CM, and is lost when macrophages are added. The ON and ON-OFF states are still observed with cancer CM (Figure 1G).

The phase portraits highlight the dynamic nature of the fibroblast-macrophage interaction, the codependence of macrophages and fibroblasts, and the strong effect of cancer condition media on these dynamics.

### Mathematical modeling infers distinct cell circuits for different contexts

The phase portrait provides a quantitative view of the dynamics that allows inferring the underlying circuits. We therefore asked whether the phase portraits (Figure 1C-E) emerge from similar or distinct underlying circuits of cell-cell interactions. To infer the cell circuits that give rise to the phase portraits, we developed a mathematical model of interacting fibroblasts (F) and macrophages (M) (Figure 2A-B), which simplifies a more complex model of biochemical reactions ^22,26^, in order to provide a minimal number of effective interaction parameters. This simplification makes it possible to infer the parameters from the data without overfitting concerns. The model has 4 parameters per cell type (Figure 2B; equations provided in Methods). Fibroblasts are removed at rate *r_F_*. Their proliferation is induced by paracrine interactions from macrophages at rate *p_MF_*, and by an autocrine loop at rate *p_FF_*. Fibroblast numbers can not exceed a carrying capacity *K_F_*, which is limited by environmental factors such as nutrients and space availability 22,29. Four analogous parameters define macrophage dynamics: removal *r_M_*, paracrine and autocrine interactions *p_FM_* and *p_MM_*, and a carrying capacity *K_M_*.

**Figure 2:**
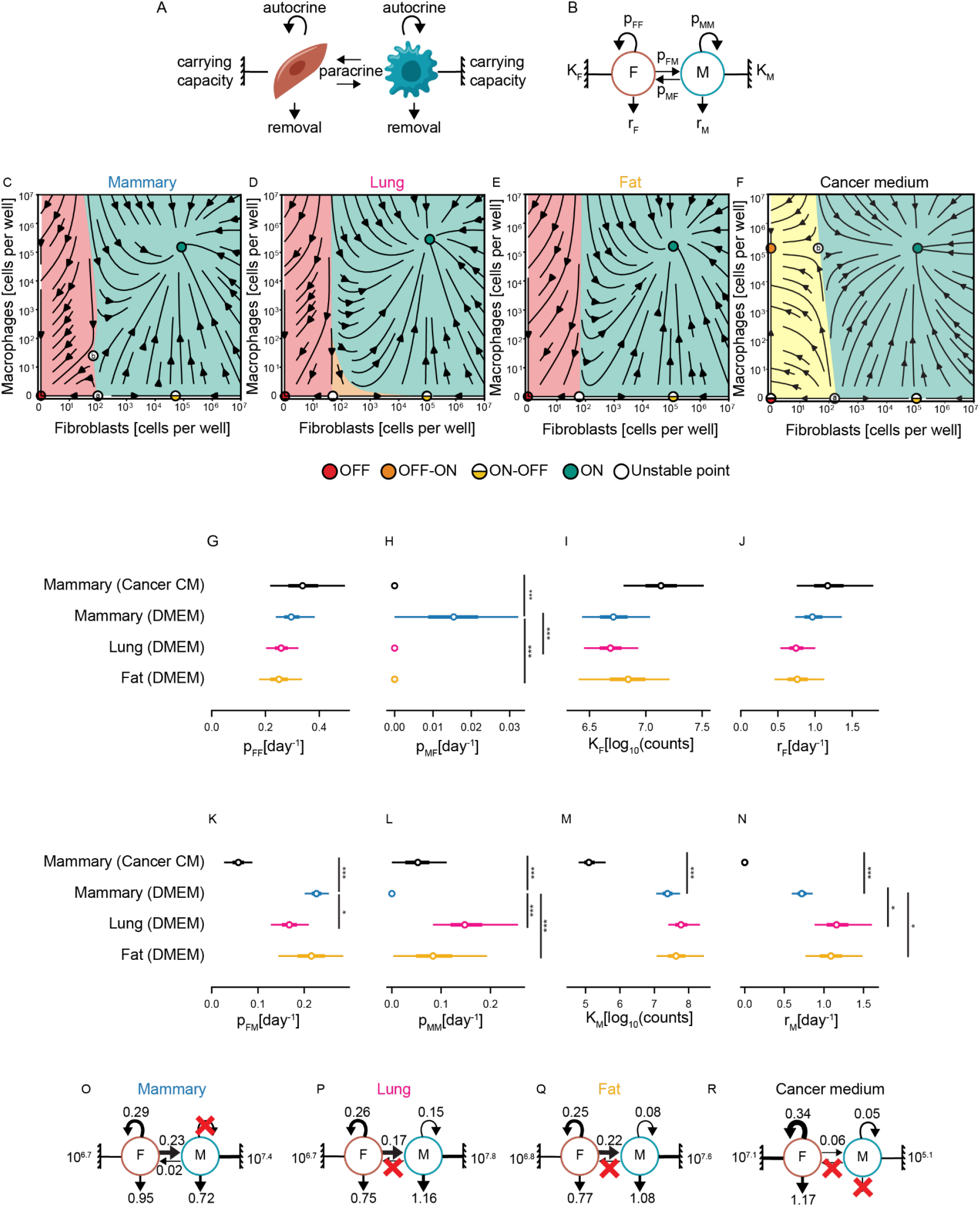
Mathematical modeling infers distinct cell circuits for different biological contexts. **A-B**. Theoretical cell circuits with the model parameters: *p_FF_* - fibroblast autocrine loop; *K_F_* - fibroblast carrying capacity; *r_F_* - fibroblast removal; and *p_MF_* - paracrine effect of macrophages on fibroblasts. Analogous parameters for the macrophages are: removal *r_M_*; paracrine and autocrine interactions *p_FM_* and *p_MM_*, respectively; and carrying capacity *K_M_*. **C-F**. Theoretical phase portraits showing dynamic interactions of macrophages with mammary, lung, fat fibroblasts, and mammary fibroblasts with cancer CM (as indicated). Basins of attraction are indicated by color: in the red area cells flow to the “OFF” state (red dot), or to the “OFF-ON” in presence of cancer CM; in the green area flow is to the “ON” state (green dot); and in the orange area cells flow to the “ON-OFF” state (half-yellow dot). **C**. Macrophages - mammary fibroblasts **D**. Macrophages - lung fibroblasts. **E**. Macrophages - fat fibroblasts. **F**. Macrophages - mammary fibroblasts with cancer CM (compared to the control mammary circuit parameters presented in Figure 2G-N). **G-N**. Calculated values for the cell circuit model parameters of mammary, lung, fat, and cancer CM. The distribution of each parameter is presented by its median (circle), interquartile range (thick line), and 95% confidence interval (CI; thin line). Parameters at zero were those removed by model selection according to information criteria. P-value was calculated by bootstrapping, *p < 0.05, ***p < 0.0005. **O-R**. Theoretical cell circuits with the mean value of each parameter for mammary, lung, fat, and cancer CM compared to the control mammary circuit parameters presented in Figure 2O (left panel).

We estimated the values of the parameters by fitting calculated to observed cell numbers at day 7 given their number at day 3, for each organ context. We tested the relevance of each parameter using standard information criteria by refitting the model when the parameter is set to zero (Figure S3A-H). The parameters discussed next are all justified by the information criteria. The model showed good fits to the experimental dynamics (Figure S3I-P) explaining 84%-93% of the variance in the data.

The models with their best-fit parameters give rise to theoretical phase portraits (Figure 2C-F, S3A-H; see Methods). These phase portraits are similar to the experimental ones, and help to fill out regions that were difficult to reach experimentally (e.g., low cell numbers). The phase portraits show the ON, ON-OFF, and OFF fixed points (Table 1, Figure 2C-F), as well as the unstable fixed points. The inferred portraits also delineate the basins of attraction for the ON and OFF states (shaded in different colors; Figure 2C-F). The border between these basins is called a separatrix.

Despite the similarity of the phase portraits in the three organ contexts, their inferred circuits were different (Figure 2G-R). Mechanistically, this suggests that the ON state is achieved differently in the mammary circuit than in the fat and lung circuits.

In all cases, fibroblasts support their own growth by means of an autocrine loop (Figure S4E-G). The ON state in the mammary circuit is achieved via a combination of the autocrine loop and a weaker paracrine interaction with the macrophages (Figure 2O). In the lung and fat, in contrast, fibroblasts grow by means of their autocrine loop without inferred paracrine support from macrophages (Figure 2P-Q).

In all organs, the macrophages are dependent on fibroblasts (Figure S4I-K). In the mammary circuit, the macrophages solely depend on paracrine signaling from fibroblasts, which pull them along to the ON state, with no inferred autocrine loop (*p_MM_* = 0). In contrast, in the lung and fat circuits, the presence of fibroblasts induces a weak inferred macrophage autocrine loop (Figure 2K-L). The enhanced proliferation rate of the macrophages is balanced by an increased removal rate (*r_M_*) in the lung and fat (Figure 2N). Lung fibroblasts also have a weaker paracrine effect on macrophages *p_FM_*, compared to mammary and fat fibroblasts (Figure 2K).

In all cases, cell growth is limited by a carrying capacity, which is similar for the three organs (Figure 2I,M). The inferred macrophage carrying capacity is 10-fold greater than the fibroblast carrying capacity, consistent with other findings ^29^ (Figure 2SG-H).

The different circuits produce differences in the phase portraits. In the mammary circuit, the model indicates a second unstable point located at low (but non-zero) numbers of fibroblasts and macrophages (Figure 2C, white dot b). From this point cells can flow to either the ON state or the OFF state. Lung and fat are missing this unstable fixed point. As a result, there is a new basin of attraction in the lung and fat phase portraits, which is missing in the mammary fat pad (Figure 2C-E, S4A-C). This basin of attraction is larger in the lung (orange region in Figure 2D).

Despite their differences, all these circuits are able to generate similar concentrations of cells at the fixed points to within a factor of two between organs (Table 1), as observed. We conclude that similar ON-states are achieved by different circuits in the three organs with lung and fat being more similar to each other than to the mammary circuit. However, all circuits have a prominent fibroblast autocrine loop, whereas macrophages depend on fibroblasts.

The theoretical phase portrait in cancer CM differs strongly from control medium. It shows a new stable OFF-ON state of macrophages (Fibroblasts are OFF, macrophages are ON; Figure 2F, orange dot), a shift of unstable fixed point *b* to higher macrophage concentrations, and a change in the OFF state from stable in the control medium to semistable in cancer CM. Although fibroblasts below a critical concentration still flow to the OFF state, addition of macrophages causes the cells to flow to the new OFF-ON state with macrophages alone (Figure 2F, Figure S4D; orange dot). Such a macrophage-only state, in which macrophages turn over and support their own growth, may resemble aspects of macrophage activation syndrome and autoinflammation ^30–32^.

This difference can be interpreted in light of the concepts of hot and cold fibrosis ^23^. In the cancer CM circuit, healing (i.e. the OFF state) is a less likely scenario, and states of hot fibrosis (ON) or autoinflammation (OFF-ON) are more likely to occur.

The new OFF-ON fixed point in which macrophages lose their dependence on fibroblasts is explained by the inferred circuit in cancer CM (Figure 2R, Figure S4L). The removal rate of macrophages is zero according to information criteria (Figure 3SH), signifying enhanced survival in the conditioned medium (Figure 2N). Cancer CM changes all macrophage parameters to allow their enhanced growth (Figure 2K-N).

**Figure 3:**
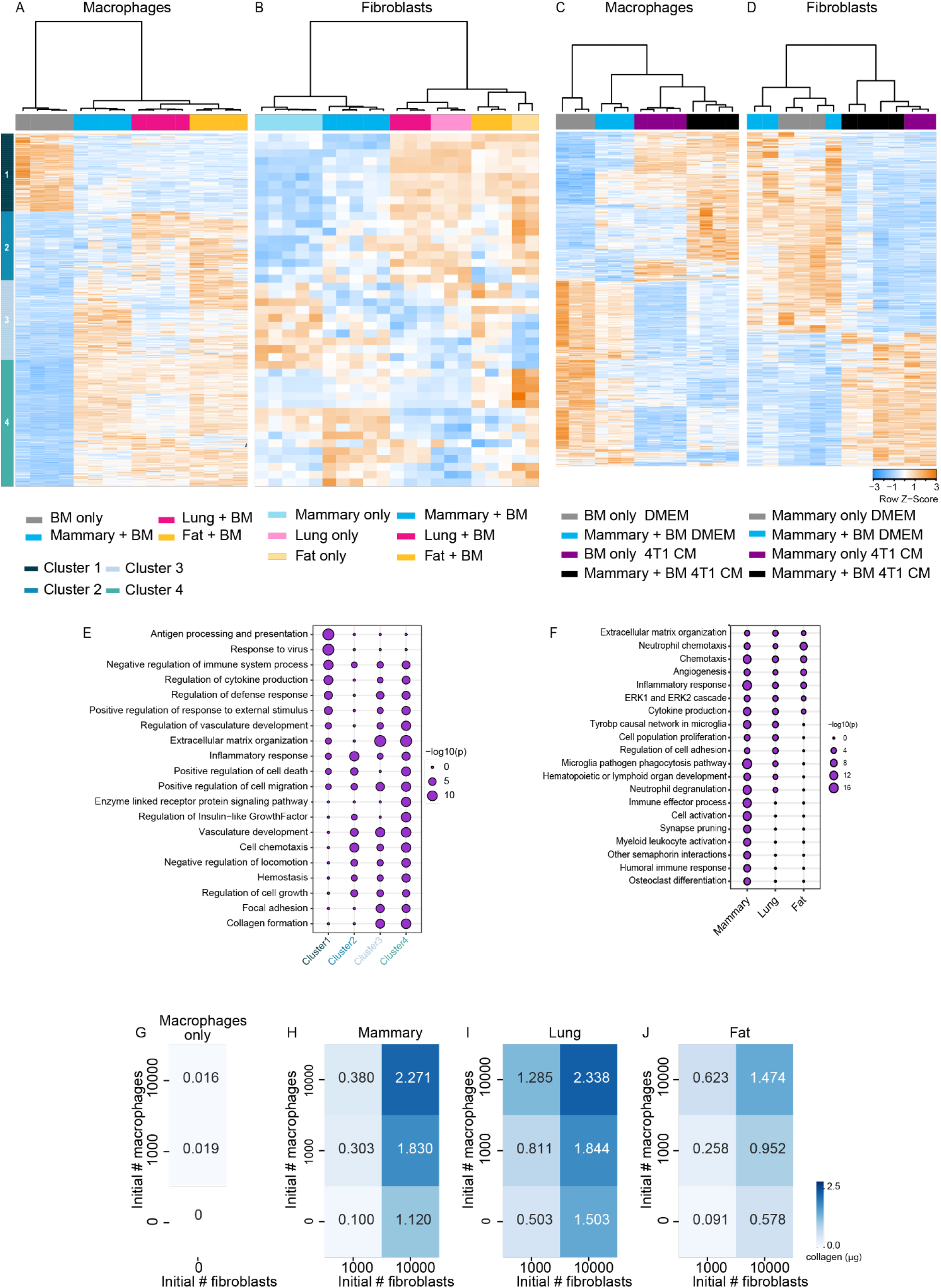
RNA sequencing supports predicted changes in macrophage and fibroblast cell circuits in different organs, and in cancer-conditioned medium. **A-D**. Heatmaps showing hierarchical clustering of differentially expressed genes (DEGs; basemean > 5; |LogFoldChange| > 1; FDR < 0.1). **A**. An additive effect model was used to compare DEGs between macrophages mono-cultured (BMDMs day0), or co-cultured with fibroblasts from different organs. Macrophages: n=4 mice. **B**. An interaction model (tissue and culture) was used to compare DEGs between fibroblasts from different organs mono-cultured (only) or co-cultured with macrophages. Mammary: n=5; lung: n=3; fat: n=2-3 mice **C**. An interaction model (medium and culture) was used to compare DEGs between macrophages mono-cultured, or co-cultured with mammary fibroblasts in DMEM *vs* cancer CM. The mono-cultured macrophages in DMEM were collected at day 0 (since they cannot maintain themselves in DMEM for 7 days), and at day 7 in cancer CM. The co-cultured macrophages were collected after 7 days of co-culture with mammary fibroblasts, in either DMEM or cancer CM. Macrophages in DMEM: n=3, macrophages in cancer CM: n=4 mice. **D**. An interaction model (medium and culture) was used to compare DEGs between fibroblasts mono-cultured (only), or co-cultured, with macrophages in cancer CM vs. DMEM. Fibroblasts in DMEM n=3, Fibroblasts in cancer CM: n=2-4 mice. **E**. Pathway analysis of the macrophage clusters from (A) was conducted using Metascape ^33^. Selected significant pathways are shown, see full list in Supplementary Table 1. **F**. Pathway analysis was performed on the differentially upregulated genes from the pairwise comparison between fibroblasts co-cultured *vs* mono-cultured (P < 0.05; FDR < 0.05; Figure S5B). Selected significant pathways are shown, see full list in Supplementary Table 2. **G-J**. Quantification of the amount of fibrillar collagen deposited as measured by Sirius Red staining after 7 days of co-culture of macrophages and fibroblasts. Macrophages only: n=4; mammary: n=3; lung: n=5; fat: n=4 biological replicates. Data are combined from at least three independent experiments. Data are presented as mean.

The cancer CM also weakens the dependency of fibroblasts on macrophages, since the inferred macrophage paracrine effect on fibroblasts is zero (Figure 2H). Fibroblast growth rate is therefore self-sustaining regardless of their seeding ratio with the macrophages (Figure S4H). Their effect on macrophages is reduced but non-zero, because the measured macrophage growth rate increases with fibroblast number in CM (Figure 2R, Figure S4L).

These analyses suggest that cancer-conditioned medium has a strong effect on the phase portrait and the inferred circuit. The two cell types become less dependent on each other. This has a profound effect on macrophage growth, since their dependence on fibroblasts in normal growth conditions is high. The effect on fibroblast growth is less profound, since they mainly depend on their autocrine loop in control conditions, rather than on the presence of macrophages. Macrophages are thus more dependent on external growth conditions - be it reciprocal signaling with fibroblasts or factors secreted to the medium by cancer cells, whereas fibroblasts are more self-sufficient and can support their own growth.

### Transcriptomic analysis reveals molecular factors underlying the circuits

Our mathematical modeling approach suggests that cancer CM strongly affects the cell circuit, and that the circuits underlying lung and fat fibroblast-macrophage interactions are more similar to each other than to the mammary fat pad. In order to understand the differences in fibroblasts between organs and the effects of cancer CM, we performed RNA-sequencing of fibroblasts and macrophages after co-culture at concentrations near the ON, ON-OFF, and OFF-ON states. Under normal growth conditions (control medium), co-culture with fibroblasts strongly affected the macrophage transcriptome, as indicated by clustering analysis (Figure 3A, first split). The genes affected by co-culture were similar between macrophages co-cultured with lung and fat fibroblasts and differed from those induced by co-culture with mammary fibroblasts, as indicated by clustering analysis (Figure 3A, second split).

Fibroblasts were not affected as strongly by co-culture with BMDMs. Their transcriptional differences correlated with organ origin, with fat and lung fibroblasts being more similar to each other than to mammary fibroblasts (Figure 3B). This supports the prediction that lung and fat circuits are indeed similar to each other and different from the mammary circuit, as well as the prediction that macrophages are affected by fibroblasts more than fibroblasts by macrophages.

To characterize these expression changes we performed pathway analysis using Metascape ^33^ (Supplementary Table 1). In macrophages, antigen processing and presentation (*H2-Aa, H2-Ab1, Cd74*) and the response to viruses (*Cxcl10, Mx1, Irf7*) were shut down upon co-culture with fibroblasts from all three organs (Figure 3A,E; Cluster 1). Co-culture with lung or fat fibroblasts led to upregulation of genes involved in inflammation and chemotaxis (*Ccr5, Bmp2, Cxcl1, Cxcl2*; Figure 3A,E; Cluster 2). The same pathways were also upregulated following co-culture with mammary fibroblasts, but the genes were different (*Ccr1, Rarres2, Ccl12*; Figure 3A,E; Cluster 3). Genes involved in ECM organization were upregulated in macrophages co-cultured with fibroblasts from all three organs (*Bgn, Serpinh1, Col3a1, Fn1, Timp1*; Figure 3A,E; Cluster 4).

Co-culture of macrophages with fibroblasts from all organs led to the upregulation of genes involved in cell growth (*Zfhx3, Slit3, Socs3*) and in positive regulation of cell death (*Apoe*, *Rhob, Bnip3*; Figure 3A,E; Cluster 4). This is consistent with the phase portrait in which the ON state is characterized by continuous turn-over of macrophages and fibroblasts, as supported by EdU measurements (Figure S1B-C).

To dissect organ- specific differences in fibroblasts, we performed pairwise comparisons of gene expression changes between fibroblasts grown alone or with macrophages (Figure S5B, Supplementary Table 2). 340 genes were differentially expressed in the mammary fat pad, 220 in the lung, and 82 in the fat. In the mammary fat pad, but not in lung and fat, co-culture with macrophages led to upregulation of genes involved in myeloid leukocyte activation (*Lat2, Trem2, Fcgr4*) and immune effector processes (*Serping1, C1qa, Fas*; Figure 3F, Supplementary Table 2). In all organs, co-culture with macrophages induced genes involved in ECM organization (*Lama2, Tnc, Loxl4*), inflammatory responses *(Adam8, C3, Cxcl10, Ccl12, Ccl6*), and chemotaxis (*Il16, Ccl12, Ccl2, Ccl20, Ccl7;* Figure 3F, Supplementary Table 2).

ECM organization pathways were induced in the RNA sequencing data in both cell types. To directly assess ECM deposition activity, we measured fibrillar collagen levels in different regions of the phase portrait (Figure 3G-J, see Methods). As expected, macrophages alone did not deposit collagen, while fibroblasts from all three organs did (Figure 3G). However, co-culture with macrophages resulted in a 2-3 fold increase in collagen deposition by fibroblasts. Maximal collagen deposition was measured near the ON state, suggesting that this state is not only the joint steady-state of the two cell types, but also the state of highest ECM production (Figure 3H-J). Taken together, the circuit analysis, transcriptional data, and ECM measurements support the notion that the ON state represents a state of chronic inflammation and hot fibrosis.

Cancer CM strongly affected gene expression in each of the cell types, and this effect was stronger than the effect of the co-culture (Figure 3C-D). Cancer CM in the ON state induced inflammation, chemotaxis, and ECM organization pathways compared to the control medium (Figure S5C-D, Supplementary Tables 3-4). This supports the prediction of the cell circuit where the cancer CM has strong effects on both cell types and weakens the dependency between them.

In summary, the transcriptomic analysis supports organ similarities (lung and fat) and differences (lung and fat *vs*. mammary), as well as the strong influence of cancer CM on macrophages and fibroblasts, highlighting the potential use of our cell-circuit approach to better understand cell-cell interactions.

### Fibroblasts depend on autocrine secretion of growth factors whereas macrophages depend on paracrine cues

The present approach identifies cell circuits and is supported by transcriptomic data. We next sought to use transcriptomic data to identify the growth factors at play in the circuits, and the relative strength of these growth factor interactions. For this purpose we employed the ICELLNET ^34^ ligand-receptor analysis framework. We focused on growth factor interactions since these are the interactions simulated in the cell circuit, and scored growth factor exchange based on the expression of ligands and receptors (Figure 4A-B, see Methods).

**Figure 4:**
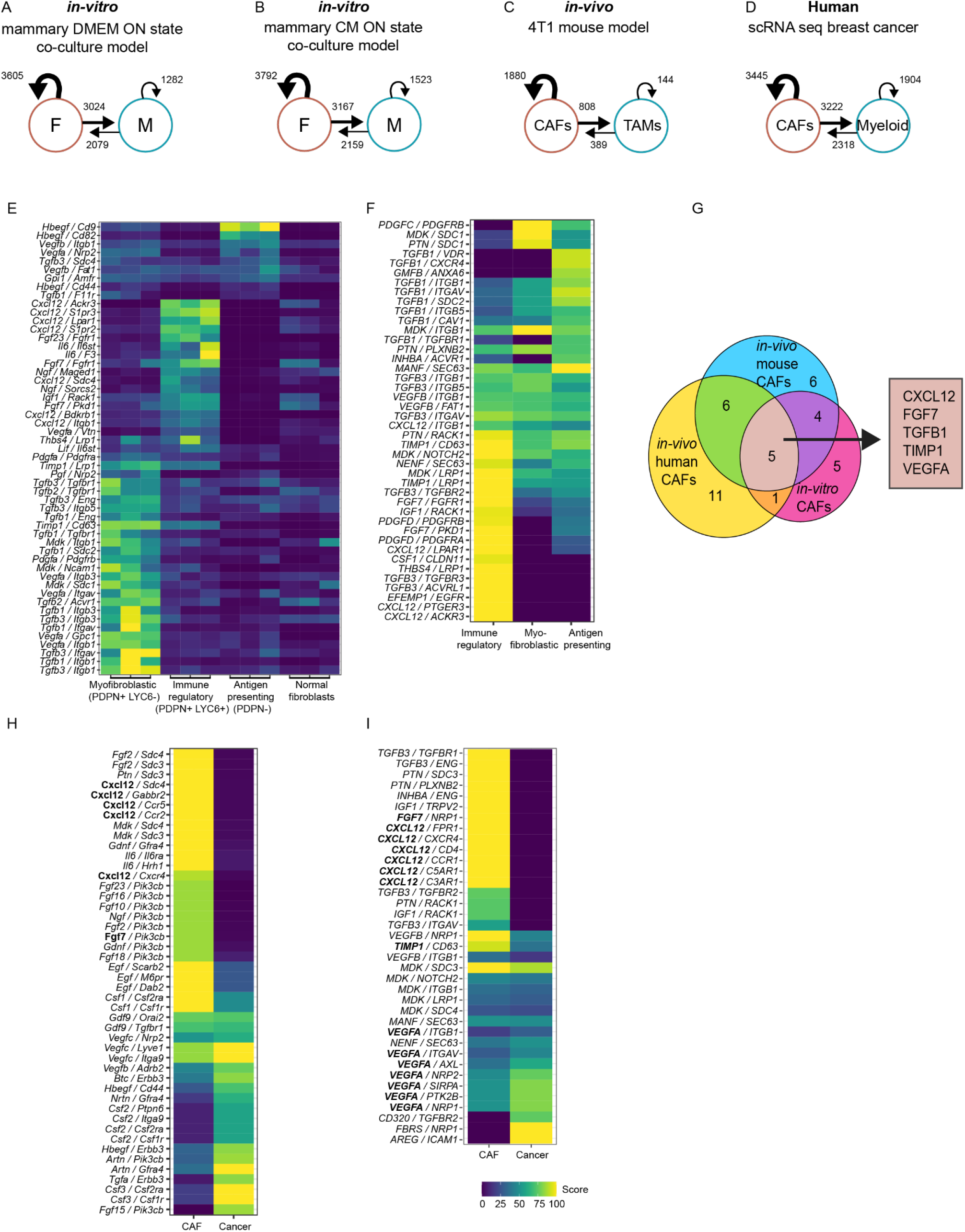
Circuits and growth factors are shared between the co-cultures and the cancer microenvironment of breast cancer from mice and human patients. **A-D**. Growth factor ligand-receptor scores for fibroblasts-macrophages based on RNA sequencing data. **A**. *In-vitro* model of mammary fibroblasts-macrophages in the ON state with DMEM. **B**. *In-vitro* model of mammary fibroblasts-macrophages in the ON state with cancer CM. **C**. *In-vivo* model of CAFs-TAMs from 4T1 tumor-bearing mice. **D**. CAFs-TAMs from scRNA of human breast cancer. **E**. The top 20 scoring growth factor ligand-receptor interactions of P_FF_ for mouse CAF populations. **F**. The top 20 scoring growth factor ligand-receptor interactions of P_FF_ for human breast CAF subpopulations. **G**. Venn diagram showing the shared growth factor ligands between fibroblasts grown *in-vitro* in cancer CM (*in-vitro* CAFs), CAF from the 4T1 model in mice (*in-vivo* mouse CAFs), and CAFs from human patients (*in-vivo* human CAFs). The gene scores were above 10, and the *in-vitro* and *in-vivo* CAF scores were at least 2-fold higher than normal fibroblasts. **H-I**. The top 20 scoring growth factor interactions of CAF and cancer ligands with TAM receptors, in mouse and human, respectively.

In both control and cancer CM, the interaction with the highest score was the autocrine loop of the fibroblasts (*p_FF_*), in which fibroblasts secrete growth factors and also express the receptor for these factors. The autocrine loop was followed in magnitude by the paracrine interaction of the fibroblasts on the macrophages (*p_FM_*), then the paracrine interaction of the macrophages on fibroblasts (*p_MF_*), and finally the autocrine loop of the macrophages (*p_MM_*). This order of strength, *p_FF_* > *p_FM_* > *p_MF_* > *p_MM_* (Figure 4A-B), is precisely the order predicted by the model for the mammary circuit (Figure 2O,R).

The specific growth factors identified in the fibroblast autocrine loop depend on the organ-context (Figure S6A-B, Supplementary Table 5). The prominent growth factors in the mammary fibroblast autocrine loop are HGF and VEGFA, whereas fat and lung are more similar to each other and include BDNF and MDK. These growth factors are known to be involved in inflammation and wound healing ^35–38^.Cancer CM affects the fibroblast autocrine growth factor interactions, independently of the presence of macrophages (S6C, Supplementary Table 5).

We conclude that the fibroblast autocrine loop is the highest scoring interaction in all contexts, in accordance with the circuit predictions.

### CAFs in human and mouse tumors show similar circuits with shared growth factors to those found in the co-culture system

To test the physiological relevance of the circuit approach beyond *in-vitro* co-cultures, we analyzed cancer associated fibroblasts (CAFs) and tumor associated macrophages (TAMs) from mouse and human breast cancer.

First, we analyzed RNA-seq data of TAMs and CAFs from a 4T1 mouse model (see Methods), and used ICELLNET ^34^ to score growth factor interactions. The CAF autocrine loop had the highest interaction score, followed by the paracrine interaction of CAFs on the TAMs, then the paracrine interaction of the TAMs on CAFs, and finally the autocrine loop of the TAMs, supporting the co-culture results (Figure 4C).

CAFs are heterogeneous and could potentially exhibit heterogeneous growth-factor interactions. To test this, we mapped the differentially upregulated growth factor interactions in each of the three main CAF subtypes - myofibroblastic (PDPN^+^LYC6^−^ pCAFs), immune regulatory (PDPN^+^LYC6^+^ pCAFs), and antigen presenting (PDPN^−^ sCAFs) that we have previously identified, and compared them to normal mammary fibroblasts ^12,18^.We found that the autocrine interactions upregulated in cancer are different between different CAF subtypes, suggesting that each CAF subtype upregulates a distinct set of biological responses in cancer (Figure S6D, Supplementary Table 5).

Next, we tested our model on human CAF and myeloid cell scRNA-seq data derived from breast cancer patients ^39^ using ICELLNET (Figure S6E-H, see Methods). Here too, the CAF autocrine loop had the highest growth factor interaction score, followed by the paracrine interactions and myeloid autocrine loop in the same order as above (Figure 4D). Furthermore, analysis of the three major CAF subpopulations: myofibroblastic (MMP11), immune regulatory (C3) and antigen presenting (HLA-DRA, CD74; Figure S6F,H), revealed distinct autocrine growth factors, supporting our findings from the 4T1 mouse model and indicating that distinct CAF subtypes engage in distinct biological autocrine interactions (Figure 4E-F, Supplementary Table 5).

Notably, the human CAF autocrine loops shared growth factors with the mouse and the co-culture autocrine loops, indicating the robustness of the present findings. The shared autocrine growth factors include *FGF7*, *TIMP1*, *CXCL12*, *TGFb1* and *VEGFA* (Figure 4G, Supplementary Table 5). TGFB1 and CXCL12 have been shown to be essential for the transition of fibroblasts to CAFs in breast cancer ^40^. The CAF autocrine growth factors mentioned above are also known to enhance the protumorigenic macrophage subsets ^1^, and are able to promote cancer invasion and proliferation ^41^. Moreover, FGF7, VEGFA, and TIMP1 promote cancer migration, angiogenesis, and remodeling of the extracellular matrix, and are known to be upregulated in breast cancer ^42–44^. Thus, these factors not only promote fibroblast proliferation but also rewire TAMs towards a protumorigenic phenotype that promote cancer progression ^45,46^. Furthermore, we found these fibroblast autocrine factors in patients with different subtypes of breast cancer (ER^+^, HER2^+^, and triple-negative). However, we did find additional unique fibroblast growth factors in patients with different breast cancer subtypes. For example, IL6 was only detected in triple-negative and HER2^+^ breast cancer patients, but not in ER^+^ patients, while FGF1,PGF and VEGFC were only detected in triple-negative breast cancer patients (Supplementary Table 5).

Finally, we analyzed the secreted paracrine growth factors expressed by the cancer cells and CAFs and affecting TAMs in human and mouse tumors. Analysis of RNA-seq data of 4T1 cells from murine tumors ^47^ and human breast cancer ^39^ revealed that cancer cells secrete factors for macrophages including CSF2 and CSF3 in mice, and AREG in patients. This provides a rationale for the OFF-ON state with macrophages alone in cancer CM (Figure 4H). The CAFs and cancer cells from both mouse and humans shared ligands that were also observed in the CAF autocrine loop, including FGF7, TIMP1, CXCL12, and VEGFA (Figure 4H-I, Supplementary Table 5). The autocrine loops for these CAFs have higher scores than the paracrine interactions from cancer cells (Figure S6I-J).

## Discussion

We present a cell circuit approach to understand the interactions between fibroblasts and macrophages in the tumor microenvironment. We defined the cell circuits using phase-portraits derived from the dynamics of *in-vitro* co-cultures. The experimental phase portraits display the dynamics for many initial conditions at once, and we developed methods to infer the underlying circuits and parameters. This allowed us to compare cell circuits from different organs and to study the effect of cancer-conditioned medium on the circuit. Fibroblasts from lung, fat, and mammary fat pad interact with co-cultured BMDMs to produce quantitatively similar fixed-point structures: an ON state with fibroblasts and macrophages in continual turnover, an ON-OFF state with fibroblasts only, and an OFF state with neither cell type. Cancer-conditioned medium profoundly changes the circuit parameters and the phase portrait, creating a new fixed point of macrophages without fibroblasts. In all contexts fibroblasts support themselves by an autocrine loop, whereas macrophages are dependent on fibroblasts or external cues supplied by the cancer cells. Transcriptomic analysis supports the circuit analysis and shows that circuits and growth factors are shared between the co-cultures and the cancer microenvironment of breast cancer from mice and human patients.

The present co-culture and modeling approach is minimal, and many physiological components are missing, including spatial structure and other cell types. Nevertheless, we find that concepts derived from the *in-vitro* system, such as the importance of the fibroblast autocrine loop, carry over to the *in-vivo* context in both mouse and human breast cancer. Moreover, some of the main growth factors are shared between the *in-vitro* and *in-vivo models*. This indicates that the phase portrait approach may be useful to study the tumor microenvironment.

The phase portrait indicates that the fibroblasts and macrophages coexist at high concentrations in an ON state with continual turnover and ECM production, as verified by EdU incorporation and collagen staining assays. This ON state is supported by an autocrine loop where fibroblasts secrete their own growth factors, and also by paracrine growth factor exchange between the cell types. The circuit analysis indicated that the fibroblast autocrine loop is the strongest of these interactions. The autocrine loop is indeed found to be the highest scoring growth factor interaction also in human and mouse breast cancer.

The shared growth factors in the CAF autocrine loop include factors previously associated with breast cancer and fibrosis. Co-inhibition of CXCL12 and TGFB1 together, was previously shown to block the rewiring of fibroblasts to CAFs in breast cancer models ^40^. The present data indicates that each of these growth factors participates in an autocrine loop of a different CAF subtype: CXCL12 in immune-regulatory CAFs, and TGFB1 in myofibroblastic CAFs. Thus, future studies may explore precision modulation of CAF subpopulations by targeting specific growth factors.

The phase portrait approach also revealed organ-specific differences in the circuits in a non-cancer context. Despite their similar fixed-points, the phase portraits for different organs are generated by distinct inferred cell-cell interaction circuits, providing an organ-specific context to the concept of cell-circuits. The organs differed in the existence of certain interactions in their inferred circuits. Fat and lung fibroblasts lacked a paracrine growth interaction from macrophages, and macrophages had an autocrine loop. In contrast, mammary fibroblasts showed paracrine growth stimulation from macrophages, and macrophages lacked an autocrine loop. This difference in inferred circuits leads to a basin of attraction seen most clearly in the lung phase portrait, that is missing in the mammary phase portrait. This new basin of attraction, in which macrophage levels decrease with time, can stabilize the ON-OFF state when macrophages are lost in a small tissue region. Thus, lung and fat may be able to stabilize a state in which fibroblasts support their own growth without macrophages, known as cold fibrosis ^23^. Our transcriptomic analysis indeed shows that lung and fat circuits are similar to each other and different from the mammary circuit.

The present organ-specific circuits may be relevant also for understanding inflammation and fibrosis. The ON state, in which high concentrations of BMDMs and fibroblasts co-exist, is known as ‘hot fibrosis’ ^23^. We find that this state is associated with high amounts of collagen deposition. The state with only fibroblasts is called ‘cold fibrosis’, and we find that it has reduced collagen deposition. In biological terms, hot fibrosis in the breast may be relevant to mastitis - a common inflammatory response of the mammary gland caused by infection or injury ^48^, whereas cold fibrosis may occur in fibrocystic disease ^48^. In the lung, a wide spectrum of interstitial diseases with varying immune involvement (from hot to cold) culminate to self perpetuating fibrosis ^49^. Fibrosis in the fat can occur in response to metabolic changes such as obesity ^50^. The inferred circuits suggest that all three organs can support hot fibrosis. These findings are supported by transcriptomic data that shows ECM remodeling and inflammatory responses induced by co-culture.

The circuit model allows one to test possible interventions *in-silico* that aim to manipulate the cell populations in the cancer microenvironment. The simulated interventions are drugs that change one of the circuit parameters, such as inhibitors of growth factor interactions - including antibodies or receptor kinase inhibitors. One may seek interventions that collapse the populations of CAFs and TAMs. Such an intervention is expected to also collapse the cancer cells due to the lack of growth factors secreted by the microenvironment ^51^ and the lack of immune inhibition offered by the microenvironment ^12,52,53^.

One favorable combination of interventions emerges from the present circuit phase portrait (Figure S6G). This combination entails inhibiting the autocrine loop of the CAFs and, at the same time, inhibiting the growth factors for macrophages secreted by the cancer cells. Inhibition of the CAF autocrine signaling loop would result in CAFs no longer supporting their own growth. The ON and ON-OFF states are both lost when the inhibition is greater than a threshold (see Methods). If this was all, the fibroblasts would collapse but the macrophages would survive, supported by the cancer paracrine growth factors. Thus, a second intervention is needed in parallel, to inhibit this paracrine support. Notably, one does not need to inhibit the interactions all the way to zero, but rather only below a certain threshold (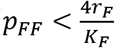, see Methods). The present data indicates that each of these inhibitions may require targeting of multiple growth factors.

The co-culture circuit approach can be generalized to other organs and cell types. One could explore, for example, different subpopulations of cancer-associated fibroblasts and macrophages, potentially together with additional cell types such as T cells and cancer cells, in order to study the circuits underlying the cancer microenvironment. The *in-vitro* system provides an accessible platform to test the effect of specific manipulations on the circuit with the aim of sculpting the cancer microenvironment towards therapeutic goals.

## Methods

### Ethics statement

All animal studies were conducted in accordance with the regulations formulated by the Institutional Animal Care and Use Committee (IACUC; protocol #05420621-2). BALB/c and C57BL/6 mice were purchased from Harlan Laboratories and maintained under specific-pathogen-free conditions at the Weizmann Institute of Science (WIS) animal facility.

### Cancer cells

4T1 murine triple negative breast cancer cells were a generous gift from the lab of Zvika Granot (HUJI, Israel). These cells were transduced to express green fluorescent protein (GFP) using the FUW-GFP vector. 4T1-GFP cells were cultured in Dulbecco’s modified Eagle’s medium (DMEM; Biological Industries, 01-052-1A) with 10% fetal bovine serum (FBS; Invitrogen) and 5% P/S (Biological Industries).

### 4T1 condition medium

4T1 cells were seeded at 1 × 10^6^ cells/ml in 10 cm plates. 24 h later (when the cells have formed a monolayer) the medium was replaced with fresh medium. 72h later, the medium was collected, filtered through a 0.22 μm strainer, and diluted with DMEM with 20% FBS, at a ratio of 1:1.

### Normal mammary fat pad and mesometrial fat fibroblasts isolation

Normal mammary fat pad and mesometrial fat fibroblasts were isolated and dissociated from the mammary fat pads or the fat tissue of two (BALB/c or C57BL/6, 8 weeks old) females per each biological replicate. organs were minced and dissociated using a gentle MACS dissociator, in the presence of an enzymatic digestion solution containing 1 mg/ml collagenase II (Merck Millipore, 234155), 1 mg/ml collagenase IV (Merck Millipore, C4-22) and 70 U/ml DNase (Invitrogen, 18047019), in DMEM. The samples were filtered through a 70 μm cell strainer into cold PBS, and cells were pelleted by centrifugation at 350*g* for 5 min at 4 °C, and resuspended in red blood cell lysis buffer (BioLegend 420301), then washed with PBS and centrifuged at 350*g* for 5 min at 4 °C. Mammary and fat fibroblasts were seeded on collagen I (Sigma-Aldrich, Cat. C3867) coated 10 cm or 6-well plates, respectively. The cells were expanded for 6 days in DMEM with 5% P/S and 10% of FBS and the media was replaced every 3 days.

### Primary Lung fibroblast isolation

Lungs of BALB/c female (8 weeks old) were excised, dissociated, minced, and incubated with enzymatic digestion solution containing 3 mg/ml collagenase A (Sigma Aldrich, 11088793001) and 70 unit/ml DNase in RPMI 1640 (Biological industries, 01-100-1A) using a gentleMACS dissociator, 30 min at 37°C. The samples were filtered through a 70-μm cell strainer into cold PBS and cells were pelleted by centrifugation at 350g for 5 min at 4 °C and resuspended in red blood cell lysis buffer, then washed with PBS and pelleted at 350g for 5 min at 4 °C. Lung fibroblasts were seeded on 10cm plates coated with collagen I. The cells were expanded for 5 days in DMEM with 5% P/S and 10% FBS, and medium was replaced after 3 days.

### Macrophage differentiation

Bone marrow-derived macrophages from BALB/c (8 weeks old) female mice were differentiated into macrophages by growth in DMEM with 10% FBS, 5% P/S and 20% L929 CM on a petri dish. The medium was replenished at day 3, and the macrophages were reseeded for the experiment on day 7.

### Macrophage-fibroblast co-culture

The fibroblasts and the macrophages were isolated and expanded for 7 days, after which the fibroblasts were trypsinized and resuspended in an ice-cold MACS buffer (PBS with 0.5% BSA). The samples were pelleted by centrifugation at 350*g* for 5 min at 4 °C, incubated with anti-EpCAM (Miltenyi, 130-105-958) and anti-CD45 (Miltenyi, 130-052-301) magnetic beads, transferred to LS columns (Miltenyi, 130-042-401), and the fibroblast-enriched, CD45/EpCAM-depleted, flow-through was collected. The macrophages were harvested with non-enzymatic cell dissociation solution (Biological Industries,03-071-1B) and washed with PBS without calcium and magnesium (PBS (−/−)). The macrophages were stained with 2 μM CFSE and seeded together with the fibroblasts in 96-well or 6-well plates precoated with collagen I. The co-cultures were grown in DMEM with 10% FBS and 5% P/S, or with 4T1-conditioned medium (performed in parallel to control media co-cultures). Every 3 days 50 μl/1ml of medium (for 96 well/6 well, respectively) were replaced with fresh medium. Macrophages and fibroblasts were seeded at different concentration ranges (0-10^5^ in 96 well and 0-5*10^6^ in 6 well), with the same combination of cell concentrations seeded in parallel onto two different plates. Plates were analyzed by Flow cytometry, one at day 3 and the other at day 7. Cell counts from 96-well plates were multiplied by 30 to scale for 6-well plates.

### Flow cytometry for cell quantification

Fibroblasts and macrophages were harvested from tissue culture plates by incubation with a non-enzymatic cell dissociation solution, washed, and transferred to round-bottom 96-well plates. The cells were then counted by flow cytometry using CFSE and anti-CD11b-Pacific blue antibody (Miltenyi, Cat.130-110-802) as positive markers for macrophages. Cells stained negatively for these markers were counted as fibroblasts. Dead cells were excluded using DRAQ7 (Biolegend, Cat. 424001). Flow cytometry was performed using CytoFlex-S (Beckman Coulter). FACS analysis was performed using flowjo software v.10.7.1.

### EdU (5-ethynyl-20 -deoxyuridine) assay

Mammary fibroblasts and macrophages were co-cultured in 96-well plates at a range of concentrations (0,1*10^3^, 1*10^4^ and 3*10^4^), and an EdU incorporation assay was performed on day 7. EdU (10 mM) was added to the cells for 2 hr, after which the cells were harvested, stained with the live/dead exclusion marker Ghost-Dye-Violet450 (TONBO, Cat.13-0863), and with anti-CD45-FITC (Miltenyi Biotec, Cat.130-110-658). EdU incorporation was detected using the Click-iT Plus EdU Flow Cytometry Assay Kit according to the manufacturer’s instructions (ThermoFisher, Cat. C10634). Samples were then acquired using a CytoFlex-S (Beckman Coulter), macrophages were gated based on positive staining for CD45, and fibroblasts were called based on negative staining for this marker. Analysis was performed with FlowJo 10.7.1.

### Collagen deposition measurement *in-vitro*

Fibroblasts and macrophages were seeded in mono-culture or co-culture (0, 1*10^3^ and 1*10^4^ cells), in 200 ul of DMEM, in collagen I pre-coated 96-well plates. Per experiment, at least two technical replicates per condition were used. Cells were left for 7 days in culture to assure confluence before performing collagen content measurement using a commercial Sirius Red collagen staining kit (Chondrex, Cat.9046), and measured by Cytation 5-Imaging Reader (Biotek).

### Cell size determination

Fibroblasts and macrophages were seeded in mono-culture at 3*10^4^ cells in 8-well slide-containing chambers (Ibidi, Cat.80826) that were pre-coated with collagen I. After 7 days, the cells were fixed in 4% paraformaldehyde (PFA) for 10 min at RT, washed twice with PBS (−/−), and stained with Dapi (to mark nuclei), and CellMask™ Deep Red plasma membrane stain (ThermoFisher, Cat.C10046), according to the manufacturer’s protocol. Images were taken with a Nikon Eclipse Ci microscope ×10 objective. Segmentation was done using Cellpose ^54^ with a Flow threshold of 0.8 and a cell probability threshold of −1 on the DAPI and CellMask channel. The cells that touched the borders were removed, and the cell sizes were quantified by QuPath ^55^ using the Cellpose segmentation.

### Bulk RNA-seq

We performed RNA-sequencing of the co-cultures at the ON state. As control, we analyzed mono-cultured fibroblasts and macrophages. Fibroblasts from different organs and BMDMs were seeded at 3×10^5^ into a pre-coated 6 well plate. The co-cultures and the mono-cultures were grown in DMEM or 4T1-conditioned medium (as described above) and were collected after 7 days. The macrophages mono-cultured were collected at day 0 since they cannot maintain themselves in control medium, and at day 7 in cancer CM. 1*10^4^ cells of fibroblasts and BMDMs were sorted using the FACSMelody instrument (BD-biosciences). All live single cells (PI negative cells after debris and doublet exclusion) were sorted. Cells staining positive for anti-CD11b-Pacific blue (Miltenyi, 130-110-802) and anti-F4/80-APC Cy7 (Biolegend. cat.123117) were sorted as macrophages, and cells staining negative for these markers were sorted as fibroblasts. The cells were collected directly into lysis/binding buffer (Life Technologies), and mRNA was isolated using Dynabeads oligo (dT) (Life Technologies). Library preparation for RNA-seq (MARS-seq) was performed as previously described ^56^. Libraries were sequenced on an Illumina NextSeq 500 machine and reads were aligned to the mouse reference genome (mm10) using STAR v.2.4.2a ^57^. Duplicate reads were filtered if they aligned to the same base and had identical UMIs. Read count was performed with HTSeq-count ^58^ in union mode, and counts were normalized using DEseq2 ^59^. Hierarchical clustering was carried out using Pearson correlation with complete linkage, and on differentially expressed genes (DEGs), which were filtered with the following parameters: basemean > 5; |log fold change| > 1; FDR < 0.1. Pathway analysis was performed using Metascape ^33^, significant pathways were determined if P < 0.05, and FDR < 0.05.

### Ligand-receptor analysis

For the ligand-receptor analysis we used the ICELLNET R package (https://github.com/soumelis-lab/ICELLNET) ^34^. We used the Nichenet dataset ^60^ and extracted the growth factor (ligand-receptor) from this dataset based on growth factor activity Gene Ontology Term GO:0008083. The scores were calculated based on the expression of genes, and as previously described ^34^. We performed this analysis on our normelazied counts from our bulk RNA-seq of co-cultured sequencing (Supplementary Tables 1-4), as described above. We also used published bulk RNA-seq data of TAMs and CAFs from a 4T1 mouse model ^61^. The normalized count genes were above 50 count in each population.

### Ligand-receptor analysis for scRNA-seq data processing and cluster annotation

We used published scRNA-seq of breast cancer patients ^39^ and published scRNA-seq of 4T1 mouse breast cancer model ^47^. We filtered cells by cutoffs of gene and unique molecular identifier count greater than 200 or lower than 10000, and a mitochondrial percentage less than 20%. We used the Seurat v.3.0.0 ^62^ method in R v.4.2.0 for data normalization, dimensionality reduction, and clustering, using default parameters. For mouse data we subgrouped myeloid, CAFs and cancer clusters by known markers that were differentially expressed between the cultures. For human data, the Normal and BRCA1 samples were removed from the analysis (BRCA1 samples had few stromal cells, and added an expression effect that we couldn’t overcome). Harmony algorithm was used to correct for patient effect ^63^, and shared nearest neighbor (SNN) modularity optimization-based clustering was then used. Cancer, Myeloid and CAF cell clusters were selected based on classic cell markers, and selected for downstream analysis (275174 cells). Matrix of normalized counts was used based on Harmony clusters. We used the same analysis including only CAF cells, and revealed several CAF subtypes (91501 cells).

### Mathematical modeling of fibroblast-macrophage circuits

The goal of the modeling was to infer the essential factors that influence the fibroblast-macrophage circuit based on the cell count data, and to compare circuits that originate from different contexts. These goals required a model with a minimal number of parameters to avoid overfitting. We therefore used steady-state assumptions for growth factor concentrations, leaving equations for the slower changes in cell numbers. We also incorporated detailed biochemical reactions ^22,26^ into a minimal number of effective interaction terms. These assumptions yielded a simple model for the rate of change of cell population, *X*, which is a balance of proliferation and removal at rates *p_X_* and *r_X_*, respectively:

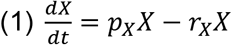

We describe the fibroblast-macrophage circuit by two such equations, one for each cell type (*X* = *F* for fibroblasts and *X* = *M* for macrophages). Fibroblasts and macrophages are removed at constant rates,*r_F_* and *r_M_*, respectively. Autocrine and paracrine interactions influence the proliferation rate, *p_X_*, of each cell population through exchange of growth factors. Proliferation is limited at high cell concentrations by resources in the medium and by contact inhibition. To account for this, we used a carrying capacity term (*K_X_*) that makes the proliferation rate decrease with growing cell population. This logistic term originates from population ecology, and was verified for fibroblasts by Zhou et al ^22^. Combining these effects yields the following equations:

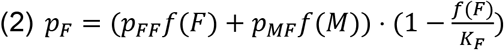

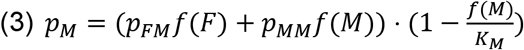

*p_FF_* and *p_MM_* are the autocrine rates and *p_MF_* and *p_FM_* are the paracrine rates. These cellular interactions depend also on the population size, *f*(*X*). Exploration of the data favored *f*(*X*) = *log*(*X* + 1), which represents a nonlinear relationship with diminishing relative effects of large cell populations. This nonlinear relationship provided better fits than a linear one, *f*(*X*) = *X*. Adding 1 inside the log is common when working with counts, in order to avoid infinity at zero cells. The function *f* resembles the saturation effect in Michaelis-Menten (MM) interactions. We chose not to use MM expressions to keep the lowest number of parameters possible, because each MM term requires an additional ‘halfway point’ parameter.

### Statistical inference

Each cell-population equation has four parameters: the rates of autocrine (*p_XX_*) and paracrine (*p_XY_*) interactions, the rate of cellular death (*r_X_*), and the carrying capacity (*K_X_*). We sought to infer these parameters from the cell count measurements.

We divided equation (1) by the population size (*X*) to obtain the per-capita growth rate, which is also the logarithmic derivative: 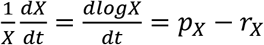. In order to fit the data we approximated the derivative as the change in cell population in an experimental time interval 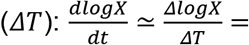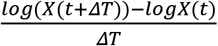. Reordering the equation gives:

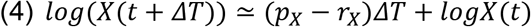

Taken together, each cell population number at day 7 (*X*_7_) can be modeled by its number and the other cell population number at day 3, *X*_3_ and *Y*_3_, where *ΔT* = 4 days:

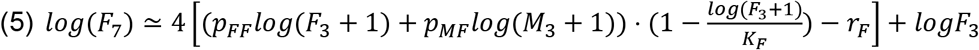

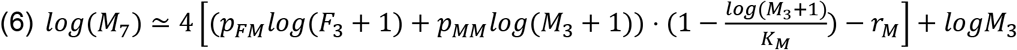

Parameters were constrained to be positive and estimated by the Trust Region Reflective (TRF) method, a nonlinear least-squares approach ^64^. We used the python implementation of this algorithm, *curve_fit* ^65^.

We tested which parameters are essential for the model fit using the Akaike Information Criterion (AIC). Parameters not justified by this criterion were set to zero.

The experimental noise of the data led to uncertainty in parameter estimations. To estimate this, we bootstrapped (resampling the data with returns) the measurements 5,000 times and inferred the parameters for each draw by the TRF algorithm. This provided a distribution for each parameter accounting for uncertainty and experimental noise.

The circuit equations with inferred parameters provided streamlines on a theoretical phase portrait. We calculated the nullclines, defined as the set of points in the phase space where there is no change in the population of one cell type (dF/dt=0 and dM/dt=0). The intersections between these nullclines provided the fixed points of the system and their basins of attraction. The net growth rate of each cell type from the model was displayed as heat maps, where growth rate changes sign at the appropriate nullcline.

### Autocrine threshold for maintaining fibroblast population

Under cancer CM, macrophages have no effect on fibroblasts growth. Therefore, the fibroblast equation is:

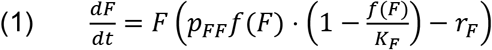

Fibroblast population crushes when 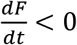. Thus:

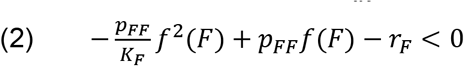

To eliminate fibroblast’s steady states and make them collapse in the whole space, inequality (2) should hold for any *F*. This happens when 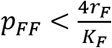.

### Statistical analysis

Statistical analysis and visualization were performed using R (Versions 3.6.0 and 4.2.0, R Foundation for Statistical Computing Vienna, Austria) and Prism 9.1.1 (Graphpad, USA). Statistical tests were performed as described in each Figure legend. In the Bulk RNA-seq, three libraries (one sample of mono-cultured fat fibroblasts and two samples of mono-cultured mammary fibroblast) were excluded due to technical problems with sequencing, as no reads were detected.

## Data availability

Bulk RNA-seq data that support the findings of this study were deposited in Gene Expression Omnibus (GEO) and can be accessed via GSE217737. All other data supporting the findings of this study are available from the corresponding author on reasonable request.

## Code availability

For the ligand-receptor analysis we used the ICELLNET R package (https://github.com/soumelis-lab/ICELLNET). We used the Seurat v.3.0.0 62 method in R v.4.2.0 to reanalyze published scRNA-seq data. Phase portraits and parameter inference of the cell circuits were calculated using Python and scipy package. Scripts and data needed to reconstruct the analysis and figures will be uploaded to github before publication.

