## Supplementary Information for "A cell circuit approach to dissect fibroblast-macrophage interactions in the tumor microenvironment"

Supplementary material – this file contains 1 Table and 6 Supplementary Figures

| | ON | ON-OFF | OFF-ON | Unstable |
|---|---|---|---|---|
| **Mammary** | $(8.9 \cdot 10^4, 2.3 \cdot 10^5)$ | $(5.5 \cdot 10^4, 0)$ | - | $(91, 0)$ $(59, 52.5)$ |
| **Lung** | $(1.1 \cdot 10^5, 2.75 \cdot 10^5)$ | $(1.1 \cdot 10^5, 0)$ | - | $(43, 0)$ |
| **Fat** | $(1.15 \cdot 10^5, 1.9 \cdot 10^5)$ | $(1.15 \cdot 10^5, 0)$ | - | $(60, 0)$ |
| **Mammary with 4T1 CM** | $(10^5, 1.3 \cdot 10^5)$ | $(10^5, 0)$ | $(0, 1.3 \cdot 10^5)$ | $(135, 0)$ $(135, 1.3 \cdot 10^5)$ |

**Table 1.** Fixed points of fibroblast-macrophage circuits. Each pair indicates the estimated cell population numbers in the fixed points by the model and data, $(F, M)$. $F$ - fibroblasts, $M$ - macrophages.

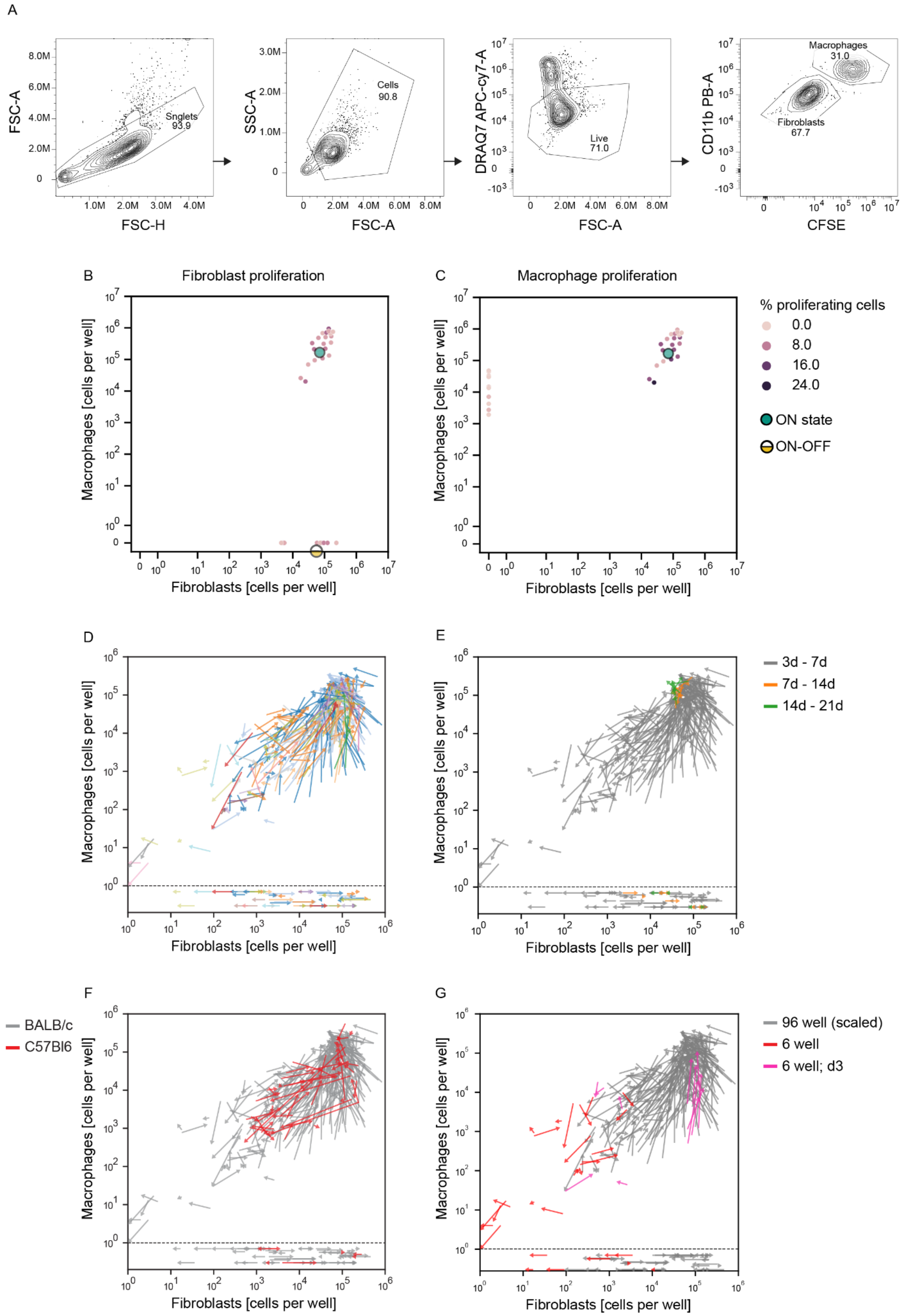
A
FSC-A
FSC-H
Snglets
93.9
8.0M
6.0M
4.0M
2.0M
0
1.0M
2.0M
3.0M
4.0M
SSC-A
FSC-A
Cells
90.8
3.0M
2.0M
1.0M
6.0M
8.0M
DRAQ7 APC-cy7-A
Live
71.0
CD11b PB-A
CFSE
Macrophages
31.0
Fibroblasts
67.7
B
Fibroblast proliferation
C
Macrophage proliferation
Macrophages [cells per well]
Fibroblasts [cells per well]
% proliferating cells
0.0
8.0
16.0
24.0
ON state
ON-OFF
D
E
3d - 7d
7d - 14d
14d - 21d
F
BALB/c
C57Bl6
G
96 well (scaled)
6 well
6 well; d3

**<u>Figure S1:</u> The phase portrait approach is robust to biological and temporal variation. A.** Flow cytometry analysis strategy: all live single cells (DRAQ7 negative cells after debris and doublet exclusion) were analyzed. Cells staining positive for CD11b and CFSE were counted as macrophages, and cells staining negative for these markers were counted as fibroblasts. Flow cytometry plots from a representative co-culture experiment are shown. **B-C.** Mammary fibroblasts and macrophages were co-cultured for 7 days after which EdU labeling was performed for 2h. Total fibroblast and macrophage numbers were counted by flow cytometry as described in **A,** and EdU+ staining was also analyzed by flow cytometry S. Total cell counts are presented as dots in the plot, and the percent of EdU+ cells is represented by shades of purple, as indicated. Data are combined from three independent experiments; n=3 biological replicates. **D-G.** Tests of robustness for the phase portrait approach to measure macrophage - mammary fibroblast dynamics *In-vitro*. **D.** Each biological replicate from the data presented in Figure 1C is presented in a different color. **E.** An experimental phase portrait comparing dynamics at different time points - co-cultures assayed at days 7 to 14 are represented by orange arrows; co-cultures assayed at days 14 to 21 are represented by green arrows. These are overlaid on the experimental phase portrait presented in Figure 1C (gray arrows). **F.** An experimental phase portrait comparing dynamics of cells from different mouse strains - measurements of cells isolated from C57BL/6 mice are shown in red, and overlaid on the experimental phase portrait of cells from BALB/c presented in Figure 1C (gray arrows). **G.** The measurements presented in Figure 1C are colored according to the following growth conditions: gray arrows represent co-cultures growing in 96 well plates, where fibroblasts and macrophages were seeded simultaneously; red arrows represent co-cultures growing in 6 well plates, where fibroblasts and macrophages were seeded simultaneously; and the pink arrows represent co-cultures growing in 6 well plates, where macrophages were added to the culture 3 days after fibroblasts were seeded.

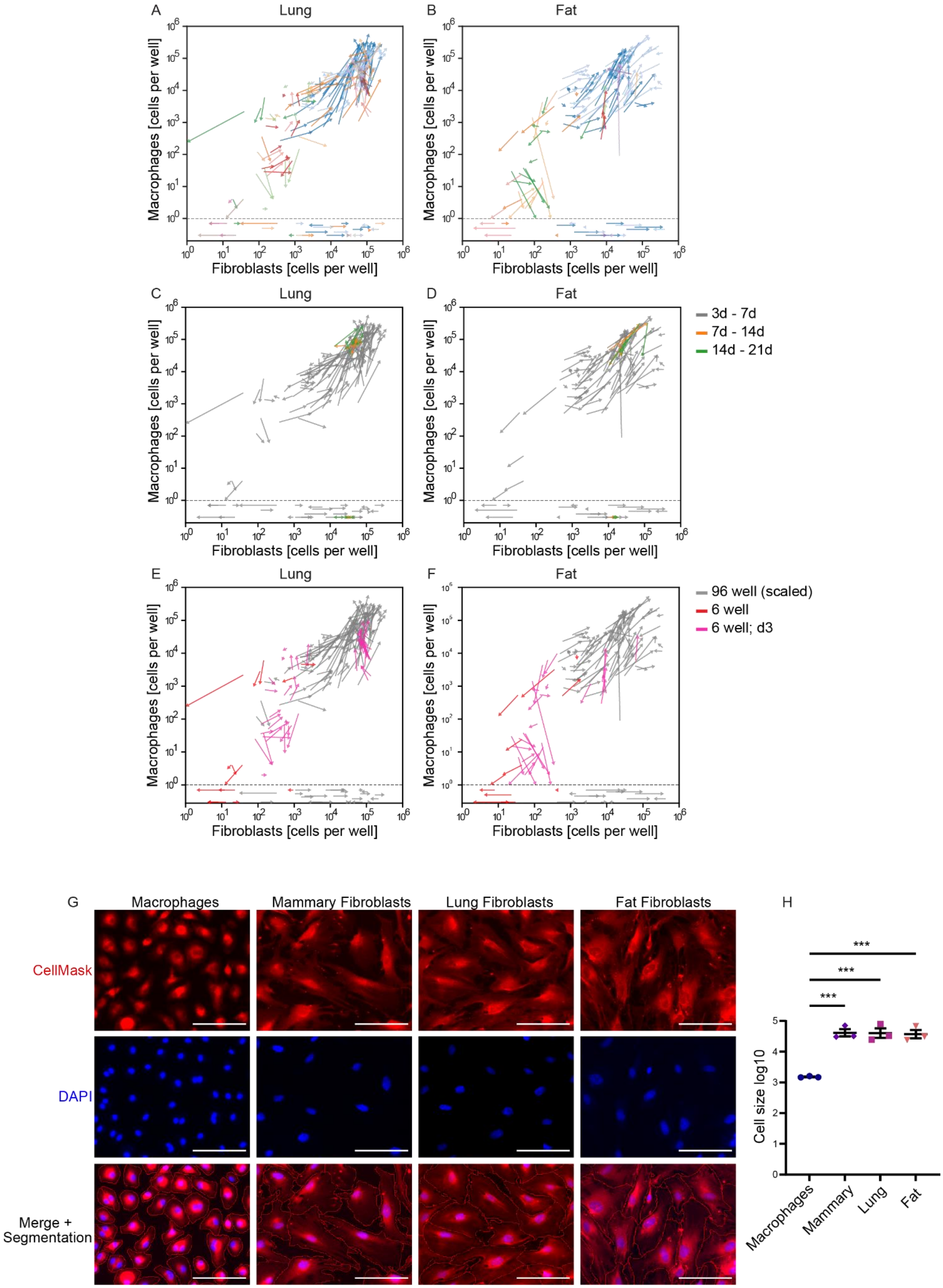
A
Lung
B
Fat
C
Lung
D
Fat
E
Lung
F
Fat
Macrophages [cells per well]
Fibroblasts [cells per well]
3d - 7d
7d - 14d
14d - 21d
96 well (scaled)
6 well
6 well; d3
G
Macrophages
Mammary Fibroblasts
Lung Fibroblasts
Fat Fibroblasts
CellMask
DAPI
Merge +
Segmentation
H
***
Cell size log10
Macrophages
Mammary
Lung
Fat

**<u>Figure S2:</u> Experimental phase portraits describing lung and fat fibroblast interactions with macrophages are robust to biological and temporal variation.** Tests of robustness for the phase portrait approach to measure macrophage - lung fibroblast dynamics are presented in **A,C,E,** and tests for macrophage - fat fibroblast dynamics are presented in **B,D,F. A-B.** Each biological replicate from the data presented in Figure 1D-E is presented in a different color. **C-D.** An experimental phase portrait comparing dynamics at different time points - co-cultures assayed at days 7 to 14 are represented by orange arrows; co-cultures assayed at days 14 to 21 are represented by green arrows. These are overlaid on the experimental phase portrait presented in Figure 1D-E (gray arrows). **E-F.** The measurements presented in Figure 1D-E are colored according to the following growth conditions: gray arrows represent co-cultures growing in 96 well plates, where fibroblasts and macrophages were seeded simultaneously; red arrows represent co-cultures growing in 6 well plates, where fibroblasts and macrophages were seeded simultaneously; and the pink arrows represent co-cultures growing in 6 well plates, where macrophages were added to the culture 3 days after fibroblasts were seeded. **G-H.** Macrophages and fibroblasts from the indicated organs were grown in monoculture for 7 days, fixed, and stained with CellMask to mark the plasma membrane and with DAPI to mark nuclei. **G.** Representative images are shown, scale bar — 67 µm. Image analysis was performed using Cellpose for cell segmentation lower panels (red line), and QuPath for cell size quantification. **H.** For each biological replicate, the average cell size was calculated from 3 images. Results are shown as mean ± SEM, n=3 biological replicates. P-value was calculated using one-way ANOVA, ***p < 0.0005.

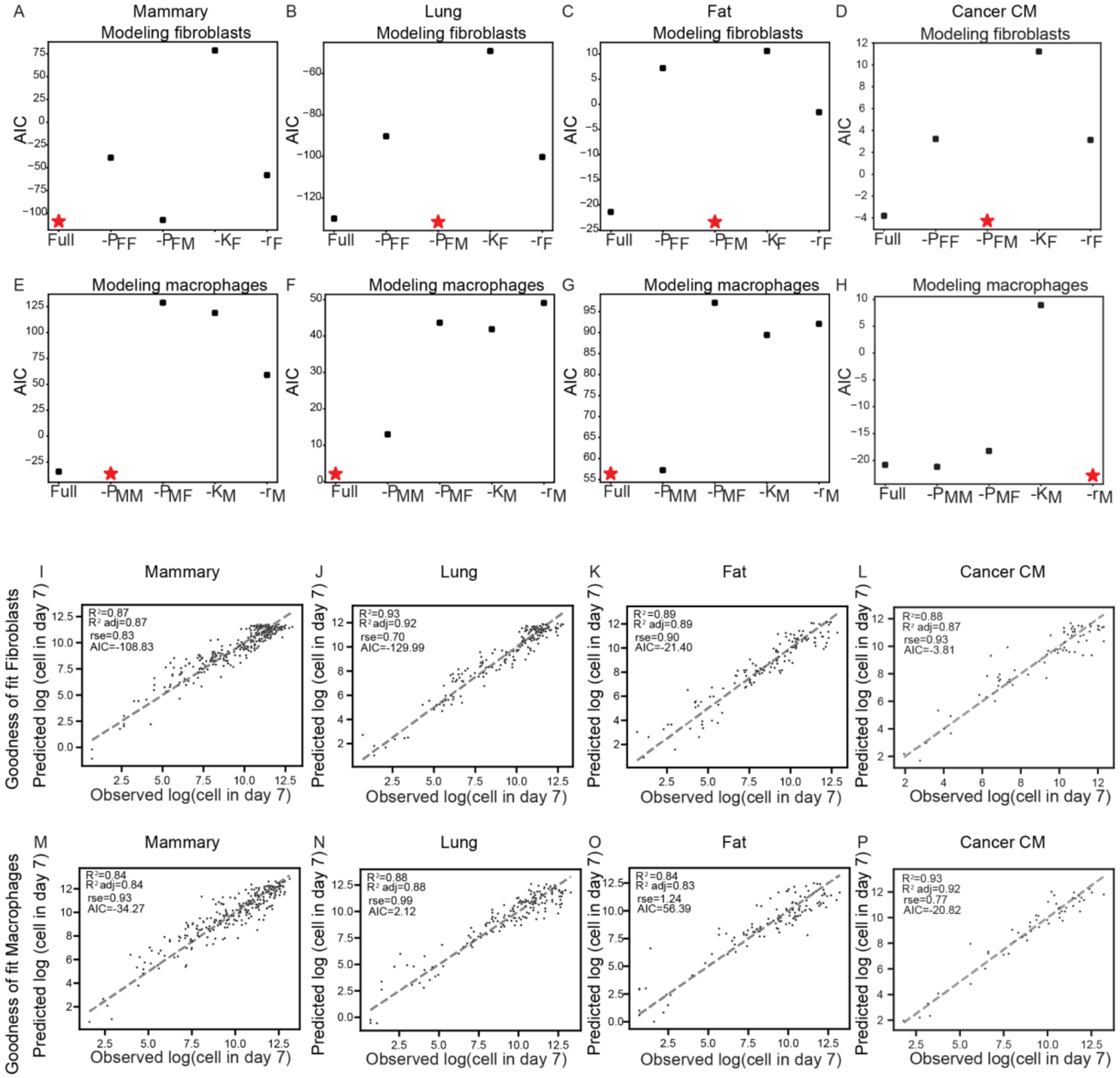


**<u>Figure S3:</u> Models chosen by Akaike Information Criterion (AIC) show satisfactory agreement with the data. A-H.** The model with the smallest Akaike Information Criterion (AIC) was chosen (red star). **I-P.** Predicted cell numbers in day 7 were plotted against the observed numbers to assess the goodness of fit of the model for **I-L.** fibroblast and **M-P.** macrophage growth. A diagonal dashed line indicates perfect fit.

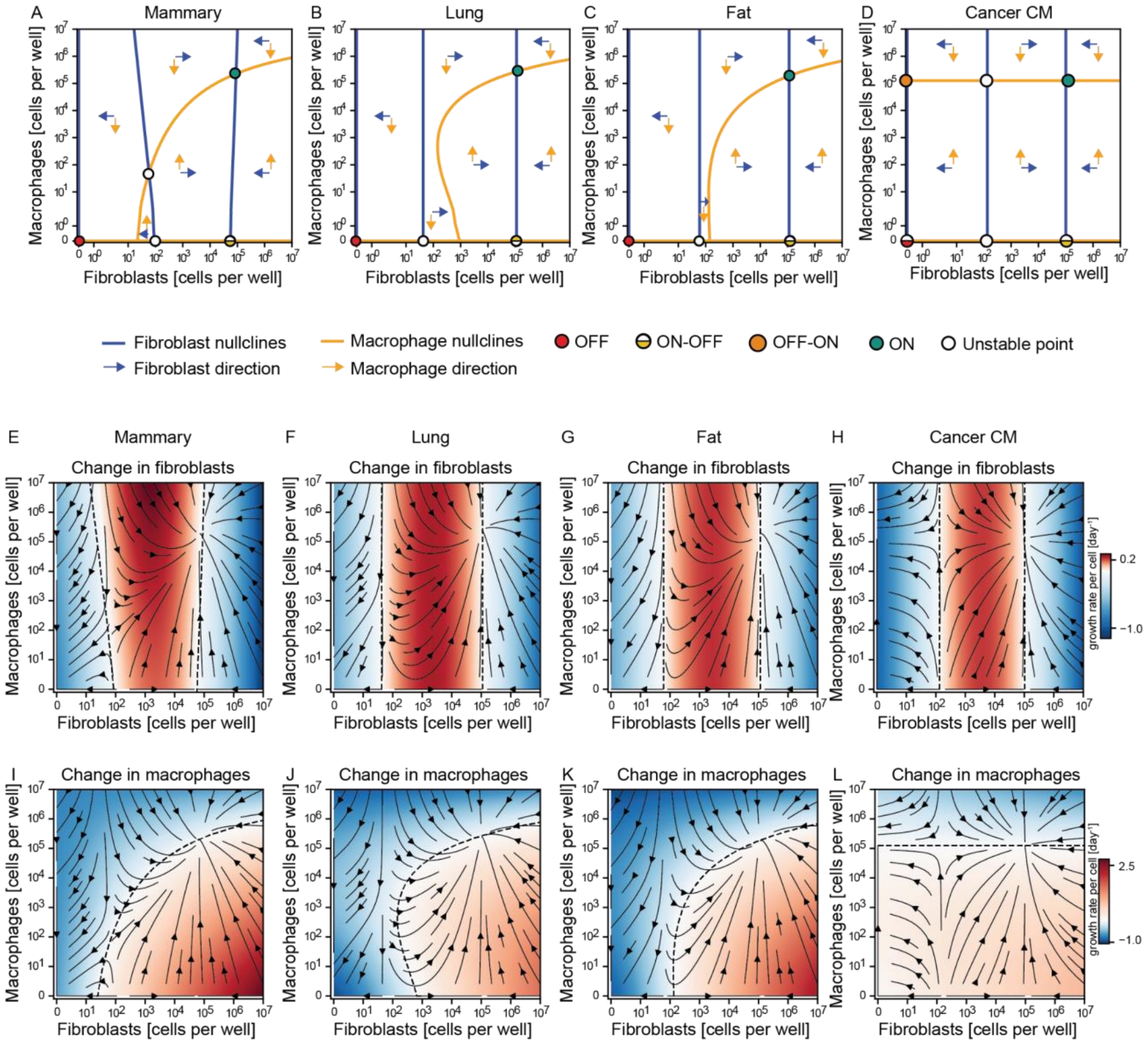


**<u>Figure S4:</u> Macrophage growth is highly dependent on fibroblasts, and this dependency decreases in cancer CM, whereas fibroblasts are more self-sufficient. A-D.** Each cell population dynamics is characterized by its inferred nullclines. The nullclines split the space into regions where the cell population increases or decreases (orange and blue arrows for macrophages and fibroblasts, respectively). The intersections of the nullclines define the fixed points of the systems (circles) and the dynamics in the vicinity of them dictate the type of the fixed point. **E-L.** Heatmaps indicating the predicted average growth rate per cell for every pair of initial conditions. Shades of red indicate that the population is growing, shades of blue indicate that the population is shrinking. Dashed lines are the nullclines of the system; along them there is no change in the cell population.

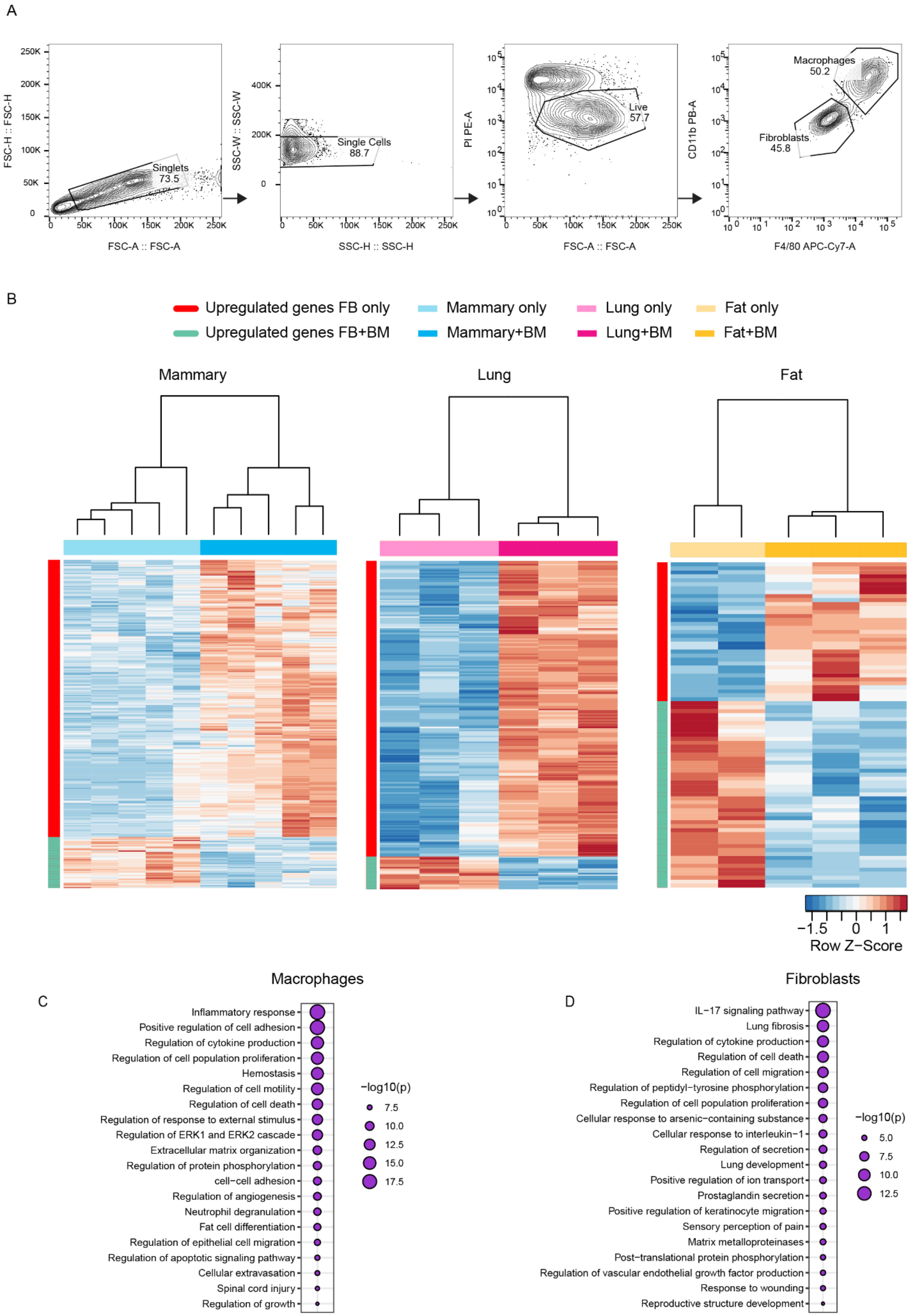
A
Singlets 73.5
Single Cells 88.7
Live 57.7
Macrophages 50.2
Fibroblasts 45.8
FSC-H :: FSC-H
FSC-A :: FSC-A
SSC-W :: SSC-W
SSC-H :: SSC-H
PI PE-A
CD11b PB-A
F4/80 APC-Cy7-A
B
Upregulated genes FB only
Upregulated genes FB+BM
Mammary only
Mammary+BM
Lung only
Lung+BM
Fat only
Fat+BM
Mammary
Lung
Fat
Row Z−Score
Macrophages
C
Inflammatory response
Positive regulation of cell adhesion
Regulation of cytokine production
Regulation of cell population proliferation
Hemostasis
Regulation of cell motility
Regulation of cell death
Regulation of response to external stimulus
Regulation of ERK1 and ERK2 cascade
Extracellular matrix organization
Regulation of protein phosphorylation
cell−cell adhesion
Regulation of angiogenesis
Neutrophil degranulation
Fat cell differentiation
Regulation of epithelial cell migration
Regulation of apoptotic signaling pathway
Cellular extravasation
Spinal cord injury
Regulation of growth
−log10(p)
Fibroblasts
D
IL−17 signaling pathway
Lung fibrosis
Regulation of cytokine production
Regulation of cell death
Regulation of cell migration
Regulation of peptidyl−tyrosine phosphorylation
Regulation of cell population proliferation
Cellular response to arsenic−containing substance
Cellular response to interleukin−1
Regulation of secretion
Lung development
Positive regulation of ion transport
Prostaglandin secretion
Positive regulation of keratinocyte migration
Sensory perception of pain
Matrix metalloproteinases
Post−translational protein phosphorylation
Regulation of vascular endothelial growth factor production
Response to wounding
Reproductive structure development
−log10(p)

**Figure S5**: **RNA sequencing reveals a transcriptional shift in fibroblasts following co-culture with macrophages. A.** FACS strategy: all live single cells (PI negative cells after debris and doublet exclusion) were sorted. Cells staining positive for CD11b and F4/80 were counted as macrophages, and cells staining negative for these markers were counted as fibroblasts. Flow cytometry plots from a representative co-culture experiment are shown. **B.** Heatmaps showing pairwise comparisons of differentially expressed genes (DEGs) between fibroblasts from the different organs mono-cultured and co-cultured with macrophages. Hierarchical clustering was carried out using Pearson correlation with complete linkage on DEGs which were filtered with the following parameters: basemean > 5; |logfoldchange| > 1 ;padj < 0.05. **C-D**. Pathway analysis was performed on DEGs from the pairwise comparison of co-cultured macrophages and mammary fibroblasts in the presence of cancer CM vs. DMEM.The analysis was conducted using Metascape [33]. Selected significant pathways (P < .05; FDR < 0.05) are shown, see full list in Supplementary Table 4-5.

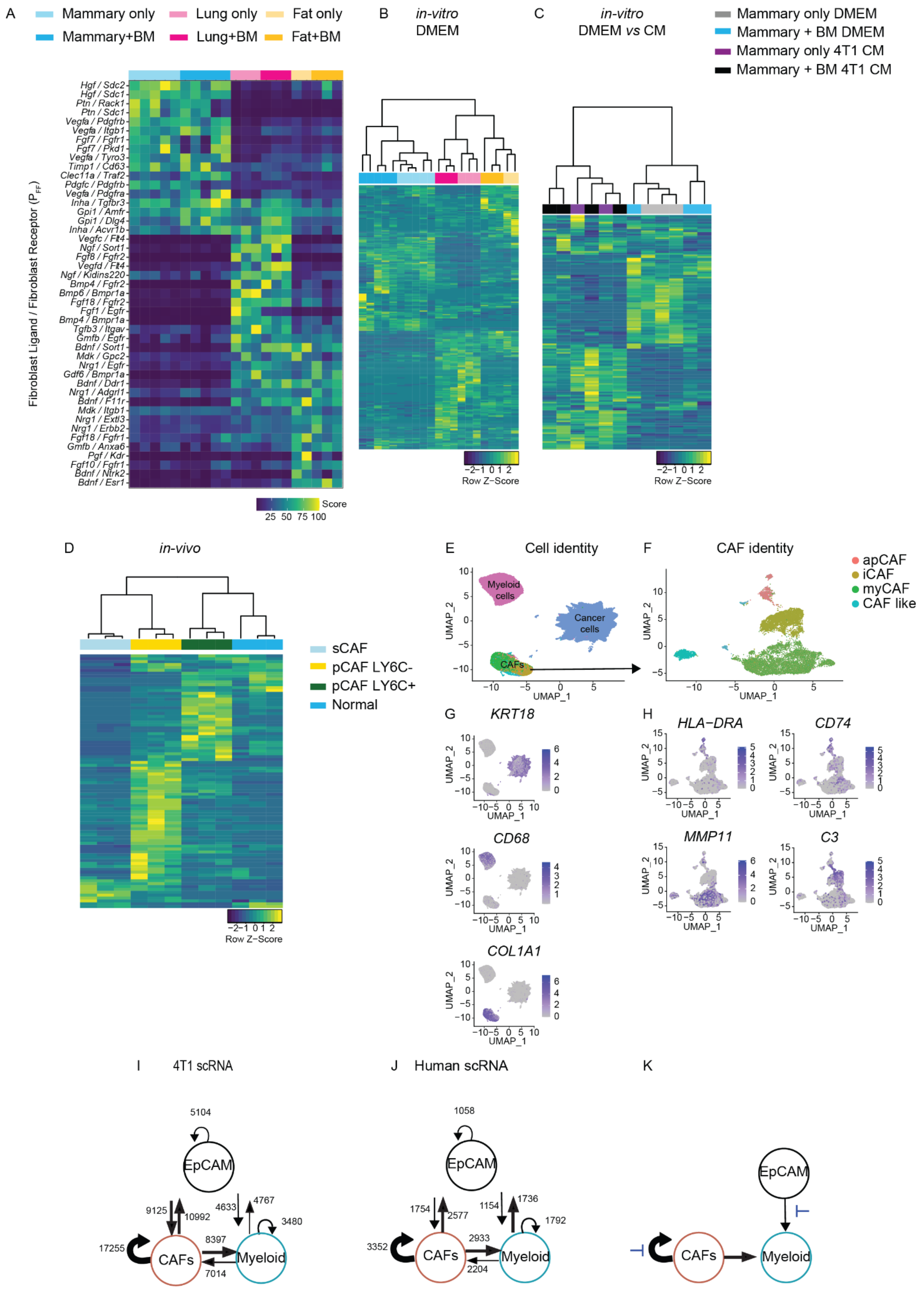
A
Mammary only
Lung only
Fat only
Mammary+BM
Lung+BM
Fat+BM
Fibroblast Ligand / Fibroblast Receptor ($P_{FF}$)
Score
25 50 75 100
B
in-vitro
DMEM
Row Z−Score
C
in-vitro
DMEM vs CM
Mammary only DMEM
Mammary + BM DMEM
Mammary only 4T1 CM
Mammary + BM 4T1 CM
D
in-vivo
sCAF
pCAF LY6C-
pCAF LY6C+
Normal
E
Cell identity
Myeloid cells
Cancer cells
CAFs
F
CAF identity
apCAF
iCAF
myCAF
CAF like
G
KRT18
CD68
COL1A1
H
HLA−DRA
CD74
MMP11
C3
UMAP_1
UMAP_2
I
4T1 scRNA
J
Human scRNA
K
EpCAM
CAFs
Myeloid

**<u>Figure S6:</u> Fibroblast autocrine growth factors depend on the organ-context and CAF subpopulation. A.** Heatmap of the top 10 scoring growth factor ligand-receptor interactions of the fibroblast autocrine loop ($P_{FF}$) of mammary, lung and fat co-cultures. **B-D.** Heatmap of $P_{FF}$ of total growth factor receptors that scored above 10 at least in one of the groups. **B.** *In-vitro*: fibroblasts co-cultured with macrophages: mammary, lung and fat. **C.** *in-vitro* : mammary fibroblasts mono-cultured and co-cultured with macrophages in DMEM vs cancer CM. **D.** *In-vivo*: CAF subpopulations (sCAF, pCAF LY6C+, pCAF LY6C-), and normal fibroblasts. Hierarchical clustering was carried out using Pearson correlation with complete linkage. **E.** UMAP visualization of cancer, CAFs and myeloid clusters after reanalysis of human scRNA seq. **F.** UMAP visualization of CAF subpopulations. **G-H.** Log-normalized expression of markers for cancer cells (*KRT18*), myeloid cells (*CD68*), CAFs (*COL1A1*), apCAF (*HLA-DR, CD74*), iCAF (*C3*), and myCAF(*MMP11*). **I-J.** Growth factor ligand-receptor scores for cancer-fibroblasts-macrophages interaction based on scRNA sequencing data **I.** 4T1 mouse circuit. **J.** Human circuit. **K.** Potential combination of interventions emerges from the present circuit phase portrait.
